# Rising water is an underappreciated driver of forest mortality

**DOI:** 10.64898/2026.09.21.753189

**Authors:** Henry Chi Hang Yeung, Tamlin M. Pavelsky, Chao Wang, Nate G. McDowell, Ryan E. Emanuel, Elsie Shenyi Liu, Emily S. Bernhardt, Xi Yang

## Abstract

Forests are known to burn and desiccate under a changing climate; less appreciated is that they also drown. Across much of the globe, a warming atmosphere is amplifying the length and intensity of both droughts and floods. Whereas massive die-offs from drought and fire are widely studied as consequences of climate change on forests, inundation-induced mortality remains poorly quantified. Using ∼1 m resolution aerial imagery and deep learning, we tracked over 260 million dead trees across the United States’ Great Lakes and ocean coastlines from 2012 to 2023. Within 10 km of the coasts, more than half of mapped mortality was concentrated in just 10% of forests, primarily in high-elevation inland wetlands and low-lying forests prone to inundation. Mortality rates doubled within a decade around expanding freshwater lakes and in inland wetlands experiencing intensifying regional-scale precipitation, while tripling in low-lying coastal forests where salinization compounded inundation in sea-level rise hotspots. Our findings identify rising waters as a widespread, underappreciated driver of forest die-off that can extend far inland, underscoring the need to reassess the impacts of climate change on the functioning, demography, and carbon-methane budgets amongst the world’s most carbon-rich forests as seas rise and hydroclimate shifts globally.

## Introduction

Widespread greening is observed in the world’s forests^1^, yet accelerating tree mortality is increasingly reported globally as climate change intensifies^2,3^. Massive forest die-offs as the result of drought and fire have been relatively well studied^4–7^, whereas tree mortality due to exacerbating inundation risk in many parts of the world has received less attention^8^. Intensifying rainfall is estimated to influence almost half of the terrestrial land surface^19^. Unlike localized hydrological disturbances from man-made infrastructure such as dams^10,11^ and artificial drainage systems^10,12,13^, climate-driven changes in precipitation can alter inundation extent and soil saturation over much broader regions^14,15^, reshaping hydrological regimes^8,14^ across both inland and coastal forests. Meanwhile, the global mean sea-level rise (SLR) rate is projected to reach 15 mm yr^−1^ by 2100 under high-emission scenarios^16^, with saltwater intrusion expected to affect nearly 80% of the world’s coastlines^17^, likely leading to extensive retreat of coastal ecosystems^18–20^. These changes are further amplified by extreme weather such as storms and drought^13–15^ and by coastal development^10,13^ that can exacerbate the inundation and salinization of low-lying areas. Importantly, these flooded and wetland forests play an outsized role in the Earth system. Although limited in extent, they are among the most carbon-dense ecosystems on Earth^22–24^, while also strongly regulating the methane cycle^25,26^, buffering millions of people from storm surges and riverine floods^27,28^, and sustaining disproportionately high biodiversity^27,28^. Accounting for inundation-driven forest loss is thus critical for comprehensively assessing climate change impacts on terrestrial ecosystems.

Emerging evidence suggests that both more frequent or longer duration periods of inundation can be associated with forest mortality^11,18,29–33^. Tree death by drowning is caused by oxygen deprivation in the roots. In waterlogged soils, gas diffusion slows and microbial oxygen demand drives the rooting zone from partial (hypoxia) toward complete (anoxia) oxygen depletion^18,34^. As oxygen declines, roots rely on inefficient anaerobic fermentation to produce energy, curtailing growth and nutrient uptake, while accelerating carbohydrate consumption^32,35^. Ultimately, prolonged exposure to anoxic soils can cause root dieback through reduced photosynthesis and phloem transport to roots. When soil saturation is accompanied by marine salinization, osmotic stress and sulfide toxicity are further compounded with oxygen stress^18,36^. However, whether these stresses eventually result in mortality depends on species-specific tolerance to hypoxia and salinity^18,37–39^, as well as local conditions such as microtopography and soil hydraulic conductivity^6,27^. Although field studies have observed forest mortality as inundation regimes change^21,37–40^, these fine-scale mortality patterns remain inherently difficult to detect with coarser-resolution satellites^41,42^, particularly in highly fragmented and heterogeneous coastal and wetland forests^29,43,44^, thus likely underestimating the true scale of inundation-associated forest loss.

Another major uncertainty is whether inundation increases mortality at the regional scale, and at what rate. Tree responses to sporadic flood pulses, such as extreme discharge or storm surge, are variable and context-dependent^11,21,37,45^, ranging from limited local impacts^21,37^ to sustained canopy loss^33,46,47^. Yet how these local responses translate into broad-scale, long-term mortality trends remains unclear. Moreover, estimates of forest sensitivity to inundation also diverge, with satellite records suggesting that forest retreat can lag SLR by multiple decades^48^, whereas field observations^21,49,50^ and models^31,51^ indicate that mortality can occur within just months to a few years of exposure to hypoxic or salinized soils. Reconciling these contrasting lines of evidence is essential for understanding future forest vulnerability to changing inundation regimes.

Here, we tracked tree mortality at the scale of individual tree crowns from 2012 to 2023 across 11 million hectares (ha) of forests within 10 km of the United States (US) coastlines, spanning the Great Lakes, Pacific, Gulf, and Atlantic coasts (Figure 1). These regions span a wide range of water-level changes, precipitation patterns, as well as salinity and elevation gradients. Using ∼1 m resolution growing-season aerial imagery from the National Agriculture Imagery Program (NAIP), we annotated 216,485 dead tree labels throughout all regions, years, and landcover types (Extended Data Fig. 1) to train deep learning models for detecting dead trees (defined as standing deadwood with white or grey crowns and associated shadows). The model ensemble achieved robust performance against manually annotated test sets in both dead tree counting (R^2^ = 0.93 and bias = −1.4% (95% CI: −8.9 to 5.7%)) and localization accuracy (F1 score = 0.76) (Extended Data Fig. 2; Methods). We then applied the ensemble to 103,356 NAIP images acquired between 2010 and 2023 to map individual tree mortality. Confident detections were matched one-by-one across years to infer their mortality timing and estimate mortality rate at the ha-scale (Extended Data Fig. 3). Using our detailed, wall-to-wall tree mortality dataset, we characterized the spatial distribution, temporal trends, and environmental drivers of mortality in three hydrologically distinct flood-prone forest systems that are especially vulnerable to rising water tables from fresh/saltwater inundation, including: (1) low-lying (< 5 m) forests along lake margins (Great Lakes), (2) inland wetland forests (> 10 m elevation), and low-lying forests along ocean coasts (Pacific, Gulf, and Atlantic; Methods).

**Figure 1.**
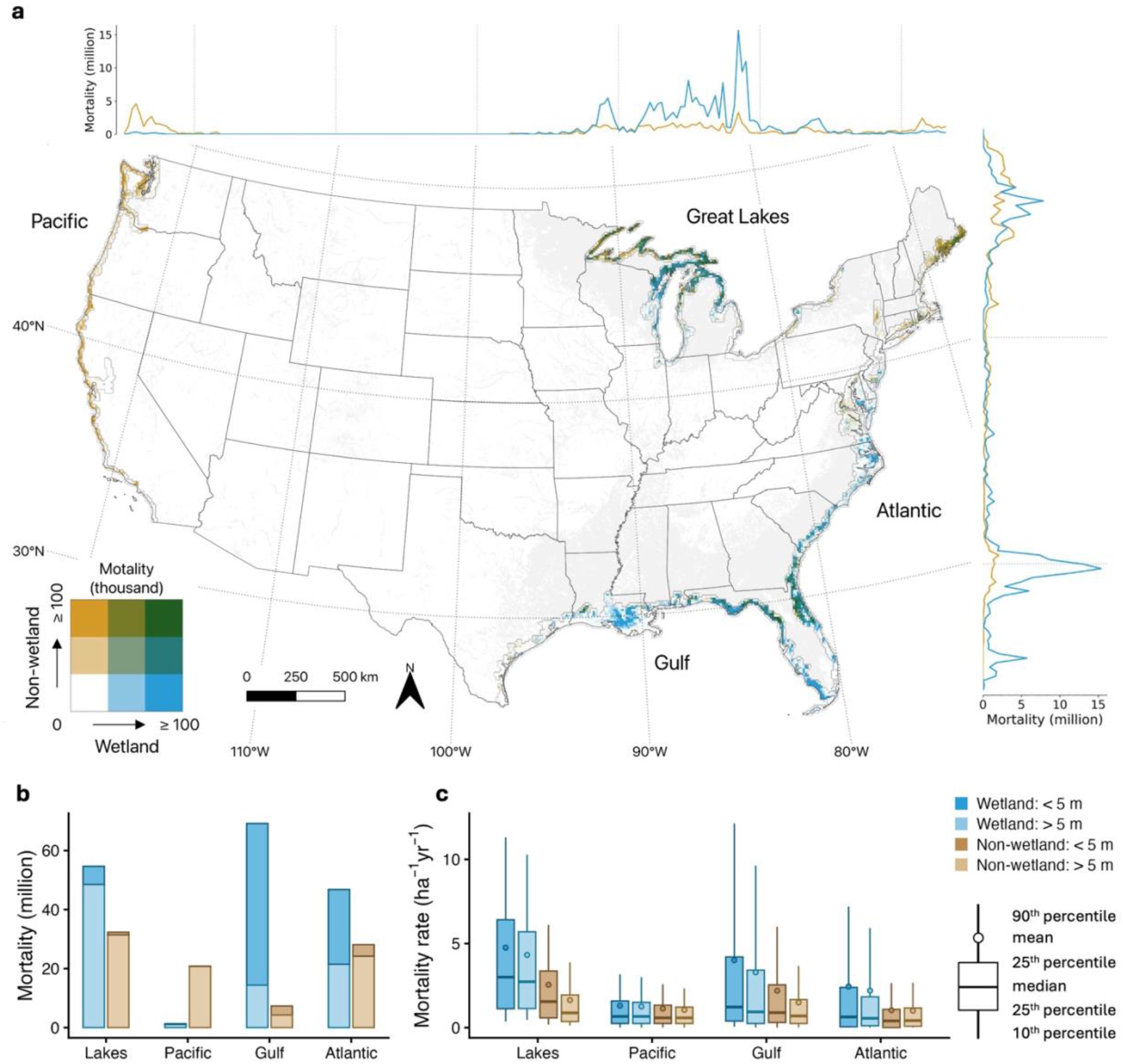
Wetland and low-lying forests contribute to substantial tree mortality. We mapped and tracked the distribution of 260 million wetland and non-wetland dead trees across the coastal US from 2012 to 2023, using ∼1 m resolution aerial images (NAIP) and deep learning models. **a**. The bivariate map shows the total number of detected dead trees (in thousands of trees per 10 km grid) in wetland and non-wetland areas during the period. Line plots show cumulative mortality (in millions) across latitude and longitude, respectively (blue is wetland; yellow is non-wetland). Basemap shows the extent of wetland forests that cover at least 1 km^2^ per 5 km grid. State boundaries from the United States Census Bureau. **b**. Cumulative mortality across four major coastal regions, colored by elevation and land cover groups. **c**. Same as **b**, but shows the mortality rate (dead trees ha^−1^ yr^−1^). Elevations are referenced to NAVD88 for oceanic coasts and the lake-specific Low Water Datum (LWD) for Great Lakes regions, respectively. Mortality associated with infestation and fire was masked using the USFS aerial detection survey datasets. Land cover from NLCD.

### Substantial tree mortality in flood-prone forests of the US

Our results showed that wetlands and low-elevation forests accounted for a disproportionate amount of tree mortality in the coastal US during 2012 – 2023. Of the 260 (95% CI: 237 to 275) million dead trees mapped, nearly 70% (180 million) occurred in these environments, despite covering only 46% of the study area (Fig. 1a, b; Extended Data Table 1). On average, wetland mortality rates (3.3 ± 5.4 (mean ± s.d.) dead trees ha^−1^ yr^−1^) were two to three times higher than that in non-wetland forests (1.2 ± 2.0 dead trees ha^−1^ yr^−1^) (Fig. 1d; Extended Data Fig. 4a, b), a pattern consistent with ground-based forest inventory estimates (Extended Data Fig. 5a). More than 60% of the mapped mortality was in the flood-exposed forests of the eastern US, including a surprisingly large amount in higher elevation (> 5 m) wetlands near the Great Lakes (19%) and northeast Atlantic regions (8%), as well as the low-lying coastal plain in the Gulf (22%) and mid-to-south Atlantic (11%) coasts (Extended Data Table 1). In contrast, mortality along the Pacific Coast was dominated by upland forests, where droughts and pests are prevalent drivers of mortality^2,7,41^. Yet, mortality rates in the Pacific Coast were less spatially concentrated (1.1 ± 1.7 dead trees ha⁻¹ yr⁻¹) than in eastern coastal systems (on average, 2.3 ± 4.3 dead trees ha⁻¹ yr⁻¹; Extended Data Fig. 4c, d) that are exposed to changing inundation patterns.

Mortality was not only widespread but also highly clustered in space. Over half of all mortality occurred within only 10% of the mapped area (Extended Data Table 1), and 76% of these hotspots were in wetlands. These hotspots (defined as areas where mortality rate exceeds 5 trees ha^−1^ yr^−1^, equivalent to 60 dead trees ha⁻¹ within the 12-year study period) accounted for 150 million dead trees across both large, contiguous wetland systems and potentially isolated inland wetlands distributed across all regions (Fig. 2; Extended Data Fig. 6). Hotspot formation can be either gradual or abrupt, depending on local conditions, and reflects multiple mechanisms, including but not limited to gradual expansion under rising lake- and sea-levels^29,52^ (Fig. 2a, f), persistent saturation in inland wetlands during climatically wet periods^33^ (Fig. 2b, c), patterned by man-made drainage networks^13^ (Fig. 2e), and abrupt post-hurricane ponding^46^ (Fig. 2f). Despite their substantial contribution to forest loss, most mortality hotspots were not captured in widely used forest loss products (Extended Data Fig. 7), suggesting that current estimates may substantially underestimate inundation-related forest mortality.

**Figure 2.**
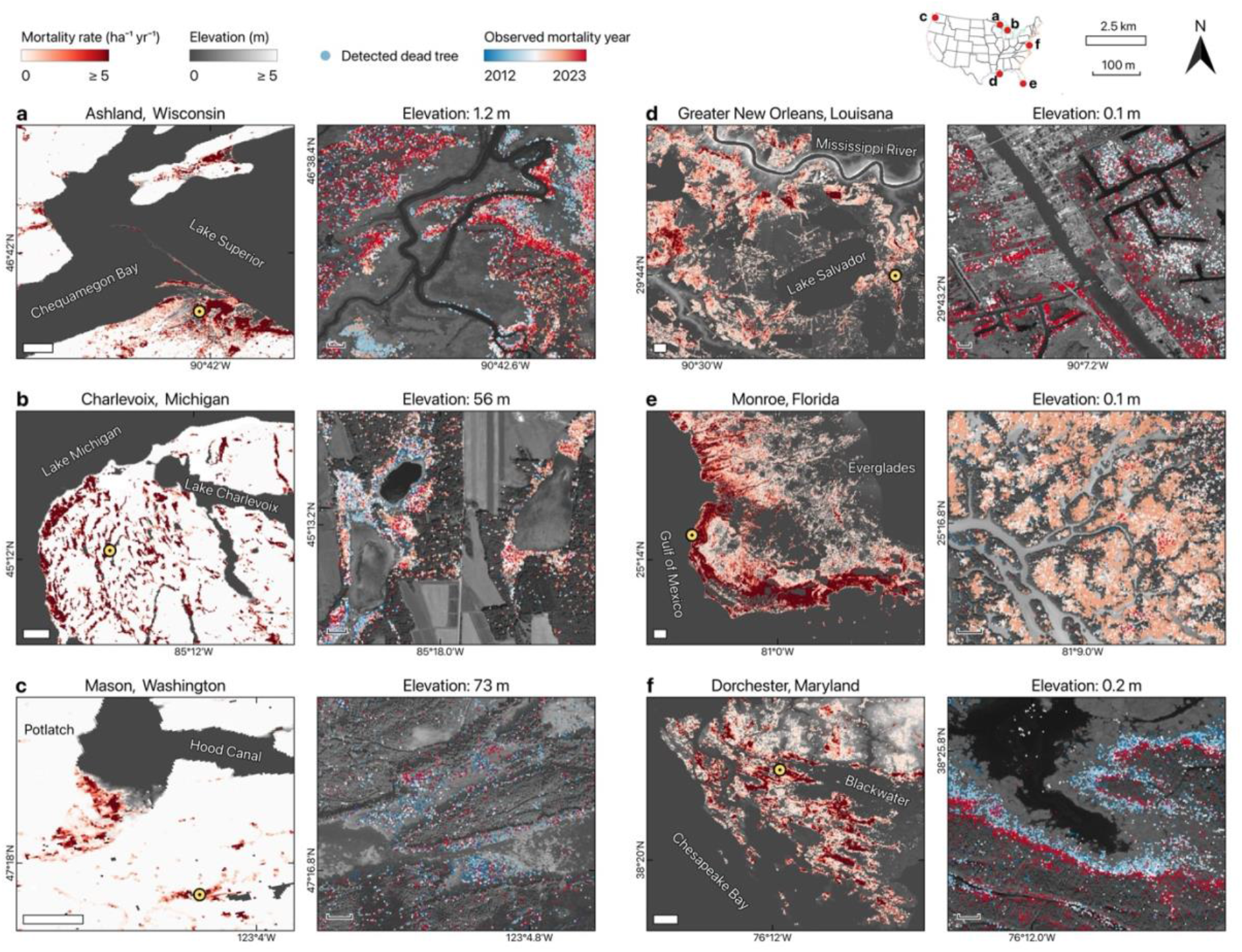
Individual tree mortality tracking reveals fine-scale die-offs in wetland forests exposed to changing inundation. **a–f**, Examples of wetland mortality hotspots across elevation gradients in the coastal US. We defined hotspots as areas with ≥ 5 dead trees ha^−1^ yr^−1^, equivalent to ≥ 60 dead trees ha⁻¹ cumulatively from 2012 to 2023, or approximating the 90th percentile of our mortality rate product (Extended Data Fig. 4). Examples include Ashland County, WI (**a**) and Charlevoix County, MI (**b**; Great Lakes), Mason County, WA (**c**; Pacific coast), Greater New Orleans, LA (**d**) and Monroe County, FL (**e**; Gulf coast), and Dorchester County, MD (**f**; Atlantic coast). Regional maps (left) show the mortality rate overlaid on elevation maps and water bodies (dark grey). These hotspots are largely undetected by existing forest loss products (Extended Data Fig. 7). Yellow circles mark the locations of high-resolution insets (right), which highlight distinct spatial mortality patterns: directional (**a**, **d**, **f**) and clustered (**b**, **c**, **e**). Colored points in insets display the observed mortality year of detected dead trees. Scale bars: 2.5 km (regional) and 100 m (insets). Representative elevations of the landscape are provided for each inset. Elevations are referenced to NAVD88 for oceanic coasts and the lake-specific Low Water Datum (LWD) for the Great Lakes regions, respectively. Mortality associated with infestation and fire was masked using aerial detection survey datasets. Basemap © ESRI.

### Increasing precipitation drives mortality in inland wetlands

Given the extensive mortality we detected near the lake margins and inland wetlands, we studied how forest ecosystems respond to purely freshwater inundation. We observed pervasive tree mortality in the periphery of rising lakes, with over 7 million trees killed along the Great Lakes (Fig. 1b, Extended Data Table 1). Between 2013 and 2020, excessive precipitation and runoff and reduced evaporation drove record-high lake levels^52,53^ (Fig. 3b), likely driving accelerated regional mortality (R = 0.68, P < 0.05; Supplementary Table 1). Trees growing on predominantly hydric soils (soil formed by waterlogging) were most vulnerable to increased precipitation frequency, with tree species exerting a secondary influence (Extended Data Fig. 8a). Frost days also contributed to mortality risk, potentially by amplifying physiological stress in northern forests^54^. Together, these results suggest that sustained waterlogging associated with rising water levels has heightened the risk of severe forest dieback along lake margins.

**Figure 3.**
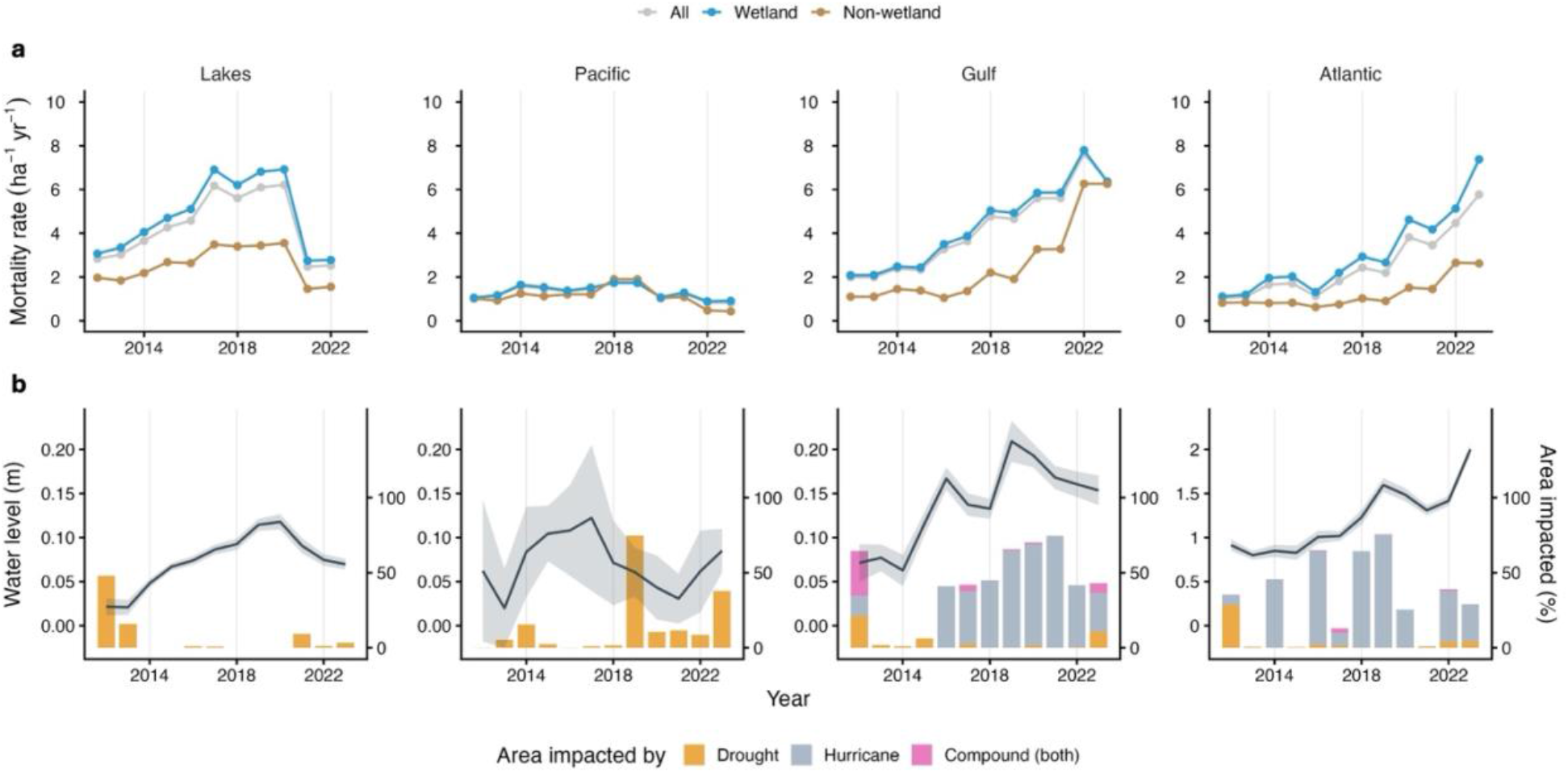
Temporal trend of mortality rate in low-lying forests along Great Lakes and ocean coasts during 2012 – 2023. Panels show temporal patterns across low-lying (< 5 m elevation) forests at freshwater (Great Lakes) and saltwater (Pacific, Gulf, and Atlantic) fronts in the US. **a**, Mean mortality rates across years in low-lying wetland and non-wetland forests. Years with less than 10% of data were not shown. **b**, Environmental stressors to low-lying forests. Lines and shaded areas show the mean water levels and 95% CI derived from tidal gauges. Bars show the percentage of low-lying forest areas impacted by drought, hurricane, or compound (both occurring in the same year), during the study period. We defined areas impacted by drought and hurricane as those with annual Standardized Precipitation Evapotranspiration Index (SPEI) < −1 and hurricane windspeed > 0 (Details in Supplementary Table 2). Water levels are referenced to the mean sea-level for oceanic coasts and to the lake-specific Low Water Datum (LWD) for the Great Lakes region.

While changing lake margins represent visible evidence of rising waters, prolonged soil saturation can also occur beneath forest canopy and well inland from lake margins due to extreme precipitation. Across the US, extensive inland wetland forests have experienced anomalously intense precipitation in recent years (Fig. 4a), coinciding with the emergence of mortality hotspots across both large and small wetlands (Fig. 4b, c). Within non-hurricane-impacted areas in our study domain, inland wetland forests exhibited mortality rates over 90% higher than matched upland forests with comparable site characteristics (Fig. 4d; Methods). Although mortality increased further when the wetlands were affected by insect or pathogen outbreaks (Supplementary Fig. 1), hydroclimatic conditions remained the dominant control on mortality risk at the regional scale (Extended Data Fig. 8b). We also found a significantly divergent response between wetland and upland forests under increasing precipitation extremes (Fig. 4e), where mortality increased sharply only in wetland forests growing on relatively poorly drained wetland soils. This sensitivity depended strongly on the precipitation regime, indicated by mean annual precipitation, to which wetland forests were acclimated (Fig. 4f; Extended Data Fig. 8b), highlighting their vulnerability to shifting hydroclimatic conditions. Similar responses were observed when we included only areas within the footprint of landfalling hurricanes (Extended Data Fig. 9), implying that episodic storm-driven rainfall may amplify mortality risk in these wetlands, although many of these affected inland wetlands were from regions with lower data quality (e.g., Florida; Methods). Collectively, these results suggest a growing role for extreme precipitation as an important compounding driver in reshaping inland wetland forests under a changing hydroclimate.

**Figure 4.**
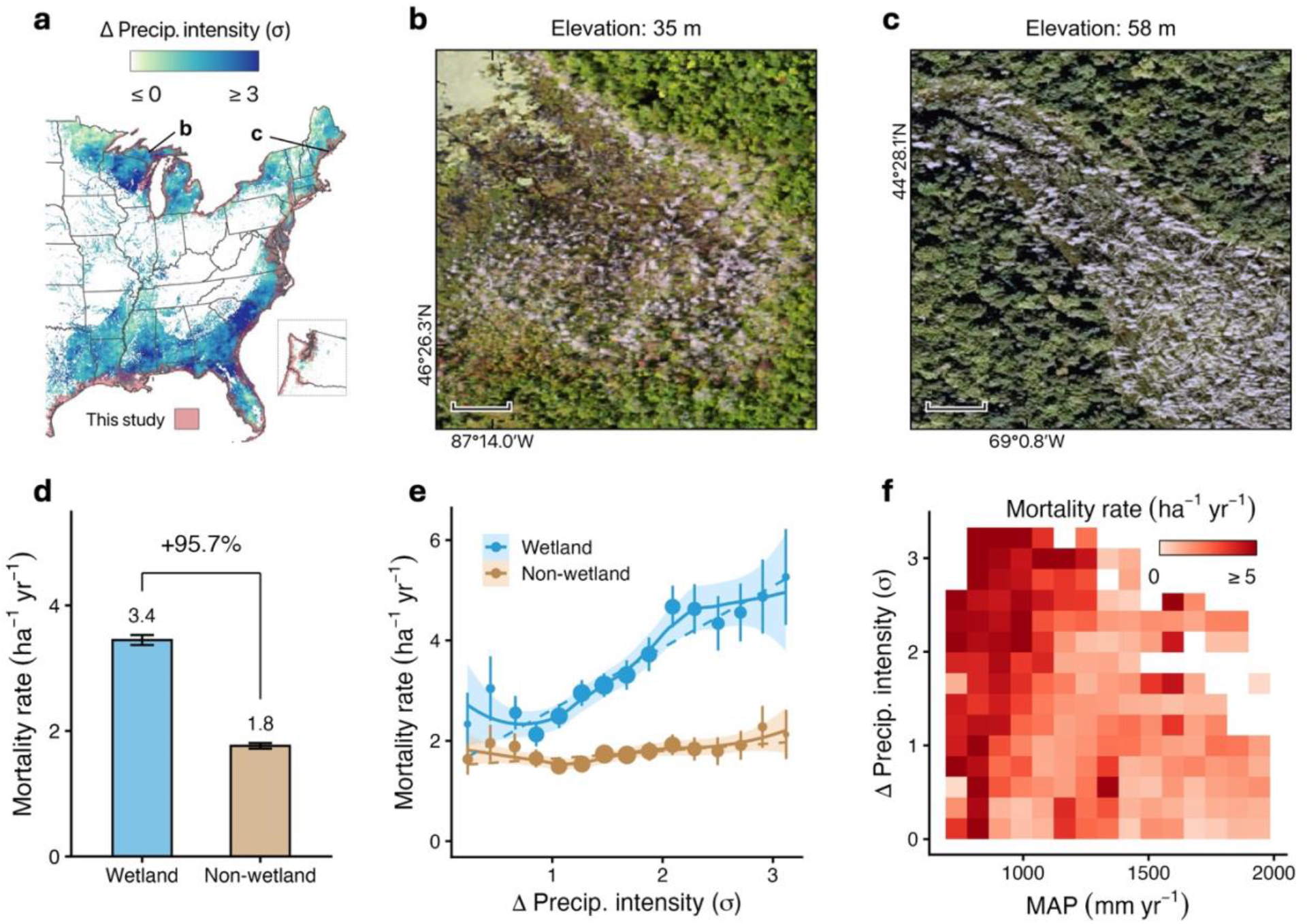
Shifting hydroclimate exacerbates mortality in inland wetlands. **a**, Standardized anomaly of precipitation intensity (2012–2023 average relative to 1980–2010 baseline) in areas with wetland forests that cover at least 1 km^2^ per 5 km grid in the eastern US. Inset: US Pacific Northwest. State boundaries from the Census Bureau. Red shading represents areas covered by this study. Examples show mortality hotspots in inland wetlands (> 10 m elevation) that emerged during the study period, including **b**, Alger County, Michigan, and **c**, Waldo County, Maine. Images from NAIP. Scale bars: 25 m. **d**, Comparison of wetland and non-wetland tree mortality rates after matching sites with similar characteristics, excluding hurricane-impacted zones (for impacted zones, see Extended Data Fig. 9). The percentage indicates the relative increase in wetland mortality compared to non-wetlands. Bar height and error bar represent the mean and 95% CI of the bootstrapped subsets (n = 25; Methods), consistent with forest inventory data even when live tree density is accounted for (Extended Data Fig. 5b). **e**, Response of wetland and non-wetland mortality rate against precipitation intensity anomaly (bin averages ± 95% CI). Point size reflects the area of forest sampled (1,000 to 25,000 ha). Dashed line is the linear regression weighted by standard error (Wetland: slope = 1.23, R^2^ = 0.85, P ≤ 0.0001; Non-wetland: slope = 0.15, R^2^ = 0.36, P = 0.02). Solid line represents a local polynomial fit for visualization, shaded with 95% CI. **f**, Wetland mortality rate against mean annual precipitation (MAP; horizontal axis) and precipitation intensity anomaly (vertical axis). Color shows the mean mortality rate of the bootstrapped samples. In **e** and **f**, the outer 2.5% of data in each tail were excluded from the distribution to ensure adequate samples in each bin.

### Accelerating mortality with rising seas

Forests susceptible to saltwater intrusion were concentrated along the Eastern US, where SLR rates exceed the global average^55,56^. We analyzed regional trends and drivers of mortality in low-lying coastal forests and found the mortality rates along the Gulf and Atlantic coasts tripled in a decade (Fig. 3a), tracking closely with relative mean sea-level (Atlantic: R = 0.87, P < 0.001; Gulf: R = 0.70, P < 0.05; Supplementary Table 1). This trend was dominated by low-lying wetlands, but the increases also occurred in forests not classified as wetlands in 2010, consistent with observations of upland-to-marsh transitions under saltwater intrusion^30,57^. Salinity of the nearest coastal water body emerged as the strongest predictor of mortality hotspot formation, highlighting the compounding role of salinization as SLR increases the frequency of saltwater exposure. In contrast, the Pacific coast—where low-lying forests are sparse—showed relatively stable mortality patterns, although localized hotspots were present in some river mouths (Extended Data Fig. 8c). Overall, forests of the Gulf and Atlantic coastal plain remained most vulnerable to future SLR, with potential for rapid, abrupt die-offs under continued climate change.

Beyond this gradual forcing, episodic disturbances also strongly modulated coastal forest mortality. Hurricane activity has significantly increased the regional mortality rate in the Gulf over the past decade (R = 0.67, P < 0.05; Supplementary Table 1). Importantly, storms were a key predictor of spatial variation in mortality in low-lying coastal forests, where storm-surge impacts are strongest, but not in inland wetlands (Extended Data Fig. 8b, c). This divergence suggests that surge-induced ponding and salinization contribute strongly to persistent root damage beyond the immediate effects of wind. Although hurricane impacts were more heterogeneous along the Atlantic coast, they still generated pronounced local mortality peaks, with mortality increasing up to tenfold within three years following major hurricanes (Supplementary Fig. 2). The magnitude and lag of these responses depend on pre-existing stressors, including pest, drought, and repeated storm exposure, which jointly regulate forest recovery dynamics^10,21,39,47^. In contrast, we did not detect a significant effect of drought alone (Supplementary Table 1, 2), potentially because drought events were relatively infrequent in the coastal plain during the study period.

Unexpectedly, higher precipitation frequency, rather than alleviating salinity stress by flushing salts from soils^10,12^, corresponded with increased low-lying forest mortality (Extended Data Fig. 8c). This counterintuitive relationship suggests that increased freshwater input may amplify forest vulnerability, potentially through interactions with SLR-driven water table rise^58,59^ and enhanced soil saturation^10,18^, amplifying root oxygen limitation under wetter conditions.

## Discussion

Inundation is an often-overlooked driver of forest die-off. By tracking individual tree mortality across flood-prone forests in the United States, we reveal that inundation-induced mortality is already pervasive across both the large and fragmented coastal plain as well as small, potentially isolated inland wetlands. Crucially, apart from rising sea-levels, an increase in precipitation variability is likely to increase tree mortality even if the mean does not change, as the effect is one-sided: our results suggest that trees will die during increasing wetting events, but they will not recover during drying periods. The scale of this mortality carries important Earth system implications. Canopy loss influences regional evapotranspiration budget^3,59,60^ and downstream hydrological processes^59,61^, while transitioning forests can alter the composition of wetland-dependent biota^62,63^ and facilitate non-native plant invasion^63,64^. Dead trees in saturated soils also contribute to long-term climate feedback by releasing carbon through decomposition and facilitating methane emissions from anaerobic soils^25,65^ for tens to hundreds of years even if the forest recovers. These findings highlight the need to incorporate flood-driven tree mortality into assessments of climate impacts on ecosystems and the terrestrial carbon budget.

Hydrological change is reshaping forests far beyond the reach of the sea. Lake expansion^15^ and increasing precipitation extremes^9,16^ are exposing inland wetlands and lake-margin forests to greater oxygen stress, while more frequent wet–dry transitions may further challenge forest recovery^52^. Across many regions, increases in both total precipitation and rainfall intensity^9^ can increase runoff and prolong soil saturation when precipitation exceeds evaporative demand. This intensification of hydrological inputs can amplify inundation in topographic depressions, driving shifts in wetland distribution^66^ and potentially promoting tree mortality in small wetlands, which play a growing role in methane emissions^25,67^. In addition, the landward expansion of rainfall footprints from landfalling storms^68^ may expose inland forests to regimes outside their historical range of variability. Inundation may make the forests more vulnerable to other stressors, including insect attacks, resulting in a compounding effect on mortality (Supplementary Fig. 1). Projections of future precipitation patterns suggest that rainfall will become more variable within the season and from year to year^16^, and a large part of high- to mid-latitude North America will see an increase in mean precipitation and extreme rainfall events^69^. However, extreme precipitation projections remain poorly constrained^70,71^, particularly at ecologically relevant scales, potentially underestimating future wetland forest vulnerability. As extreme rainfall events continue to intensify, inundation of inland forests is likely to become an increasingly important contributor to global forest change.

Our wall-to-wall mortality mapping also suggests that coastal forests might be more sensitive to saltwater intrusion than previously recognized. The strong coupling between regional mortality rates and SLR, together with field observations documenting tree death within months to years of salinization^39,49^, implies that forest transitions can occur within years rather than multidecadal timescales. Whether such dynamics facilitate saltwater intrusion will depend on freshwater inputs and groundwater levels. Whether salinity or inundation stress ultimately dominates remains unclear and will depend on freshwater inputs and groundwater levels, with extreme rainfall potentially exacerbating oxygen stress. Although many coastal ecosystems are projected to collapse when SLR rates exceed 7 mm yr⁻¹ ^56^, this rate has already been surpassed in some regions, such as the US coastal plain, owing to interannual climate variability^56^. Wetland elevations can keep pace with SLR to a certain degree and decelerate their retreat^59^, but accelerating global SLR projected throughout this century^16^, compounded by storms, droughts, and anthropogenic modifications of coastal landscapes^10,12,72^, may drive substantial losses of coastal forests sooner than current projections suggest.

Our approach also highlights key challenges and opportunities for monitoring flooded forests. We infer inundation impacts from environmental proxies because robust detection of under-canopy inundation remains difficult in dense, high-biomass forests^73^. This limitation is particularly important for heterogeneous flooded forests, which are therefore often underrepresented in regional and global forest loss products^74,75^, despite accounting for substantial mortality in our analysis. Emerging low-frequency microwave missions, including NASA-ISRO Synthetic Aperture Radar (NISAR)^76^ and ESA BIOMASS^77^, may substantially improve the detection of standing water beneath forest canopies^73^ and enable more comprehensive assessments of wetland vulnerability and distribution shifts across spatial scales. In addition, the dead-tree-tracking confidence and temporal resolution of our mortality record are constrained by the availability of high-quality very-high-resolution imagery, hindering our ability to quantify forest resilience against inundation stress, particularly during stabilizing periods with lower water levels and less extreme weather. While recovery in mortality hotspots is sometimes observed in our map (Supplementary Fig. 3), potentially linked to carbohydrate reserves within trees^32,49^, it should be interpreted with caution given these data limitations. Our approach likely misses some small trees (e.g., crown diameter < 1 m) that are not detectable or understory trees that are occluded; thus, our dead-tree count should be viewed as a conservative estimate. Nevertheless, comparison with forest inventory data suggests that our results broadly agree with ground observations (Extended Data Fig. 5). Increasing availability of high-resolution satellite data can improve monitoring of individual tree health. Interpreting potential drivers presents additional complexity. Notably, mortality is higher in frequently infested areas than under high precipitation variability, despite infestation showing little regional importance (Extended Data Fig. 8). This mismatch likely reflects both uncertainty in infestation maps^78^, as well as potentially more localized extent of infestation relative to a change in inundation pattern at regional scale. Integrating field and experimental measurements, long-term satellite observations, and high-resolution mortality mapping could help resolve these dynamics and improve understanding of forest dynamics to hydrological extremes.

Fire leaves a scar that satellites can see from space; drowning does not. That asymmetry has let inundation-driven forest mortality go largely unaccounted for in assessments of climate impacts on the world’s forests, even as it already rivals fire and drought in scale across US coastal and inland wetlands. As seas rise, storms intensify, and rainfall regimes shift, this hidden driver of forest death is poised to grow. Bringing drowning into view alongside burning will be essential for accurately forecasting the fate of the world’s most carbon-dense forests.

## Methods

### Overview

To assess inundation impact on tree mortality, we mapped individual dead trees across the coastal US (within 10 km of the coastline) from 2010 to 2023 using ∼1 m resolution aerial imagery and a deep learning ensemble. Using parallel computing, we tracked detected dead trees to identify new mortality for each observation year and produced biennial mortality rate maps at 100 m resolution for 2012–2023. With these high-resolution mortality maps, we analyzed the spatial and temporal distribution of mortality rates across three flood-prone systems — low-lying (< 5 m elevation) forests along lake margins (Great Lakes), inland wetlands (> 10 m elevation), and low-lying forests along ocean coasts (Pacific, Gulf, and Atlantic). We used 5 m as the upper elevation threshold for low-lying coastal forests because it is commonly used to assess coastal ecosystem vulnerability to rising waters and storm surge^29,48,79,80^. We also kept this threshold consistent for the Great Lakes region. Additionally, since extreme storm-surge events have historically reached elevations of up to 8.5 m^81^, we excluded forests between 5 and 10 m from inland wetland analysis to minimize the potential influence of saltwater. We then assessed the dominant environmental predictors that explain the formation of mortality hotspots using random forest models trained on geophysical, climatic, and biotic predictors. Finally, we assessed the response of inland wetlands to extreme precipitation and compared their mortality rates to paired upland forests with similar characteristics.

### Aerial imagery

We acquired aerial imagery from the National Agricultural Imagery Program (NAIP) to map individual dead trees across the coastal US. NAIP images are collected throughout the US during the growing season every 2 to 3 years with 4 spectral bands (red, green, blue, and near-infrared)^82^. All images have been orthorectified and atmospherically corrected by the vendors. Georeferencing accuracy improved from ±6 m (95% confidence) prior to 2017 to ±4 m thereafter. We used the more conservative bound (i.e., ±6 m) as the spatial matching radius in all subsequent tree-tracking analyses. To standardize the resolution of NAIP images acquired across states and years, which varies from 0.3 – 1 m, all images were bilinearly resampled to 1 m resolution. We downloaded 103,356 images (25 TB) collected from 2010 to 2023 across 27 states spanning all US coasts using Google Earth Engine (GEE).

### Coastline, elevation, land cover, and environmental data

We define the study domain as areas within 10 km of the coastline based on Euclidean distances from the NOAA medium-resolution shoreline^83^. We applied the USGS annual National Land Cover Dataset (NLCD) to mask agricultural fields and urban areas each year, and used the 2010 land cover as the baseline for wetland and non-wetland forest classification. Our product included areas with at least 10% tree cover according to the global tree cover map in 2010^84^, but retained only those classified as upland forests or woody wetlands for all subsequent analyses. Elevation was derived from the USGS 3D Elevation Program light detection and ranging (LiDAR) digital elevation model (DEM)^85^. This LiDAR DEM is referenced to the North American Vertical Datum of 1988 (NAVD88) vertical datum, which is associated with local mean sea-level and has a vertical accuracy of ± 0.82 m. The elevation for each lake was then referenced to a stationary water level baseline using the Low Water Datum (LWD)^86^. The details and processing of all environmental variables, including the water levels, hurricanes, droughts, and precipitation indices, are described in Supplementary Table 2.

### Mapping individual tree mortality with deep learning

To reliably map individual standing dead trees, we utilized a deep-learning architecture based on Gaussian heatmap regression^87^, originally designed for high-density object counting. This approach was chosen over crown segmentation since it requires only point annotations for training, which drastically reduces labeling effort across a domain of this geographic extent^88^. In total, we compiled 226,078 dead tree labels spanning all coastal regions from 2010 – 2023 to train and evaluate our models. The modeling framework was implemented in Python (3.12) using TensorFlow.

Model training proceeded in two primary phases. A base network was first trained on 132,490 labels from the Atlantic coast, covering 16,610 ha^29^, then fine-tuned on newly generated labels from all year-regions of the US. Since mortality hotspots are rare events, we identified them through extensive searches in the literature^18,59^, news articles, and Google Earth Pro imagery, and applied stratified random sampling for the remaining non-hotspot locations to capture diverse coastal land covers (e.g., deciduous and evergreen forests, wetland, shrubland, grassland, agricultural, and urban). Non-forested land covers were included in the training samples to minimize the potential propagation of dead tree detection errors during spatial aggregation. Next, we extracted image patches (either 256 × 256 or 512 × 512 pixels) for these selected locations and applied a semi-supervised labeling framework to reduce labor for dead-tree labeling. We manually annotated more than 50,000 labels in these image patches, verified by five trained specialists. The base model was subsequently applied to generate pseudo-labels, where high-confidence labels were manually inspected before incorporation into an iterative retraining cycle, and until the model outputs were visually satisfactory. This process yielded a dataset of 83,995 labels across 5,720 images, covering 11,945 ha of all year-regions for model training (Extended Data Fig. 1). Thus, the total number of labels we created is 216,485.

The network was modified to produce three output components: a Gaussian heatmap estimating the detection confidence of individual dead trees^89^, a scale map adjusting Gaussian spread to accommodate crown size variation^88^ – thereby enhancing accuracy in dense forests – and an attention map directing the model’s area of focus^87–89^. We added location embeddings (i.e., latitude, longitude) to improve generalizability across the climatically diverse study domain. The model was optimized using a weighted scale-adaptive Gaussian loss, parameterized by α and β to balance commission and omission errors, and γ, a focal component that can down-weight hard samples. Because training labels were annotated conservatively, smaller α and γ were chosen to reduce false-negative predictions^2^. Following ref. ^87^ and hyperparameter tuning, we set α = 0.3, β = 0.7, and γ = 0.8 for the loss function, and initialized the Gaussian kernel size as 3 standard deviations and a minimum scale-map value of 0.5 m. The model was trained on 256 × 256 patches at batch size 12 using the Adam optimizer^90^, with learning rate initialized at 0.001, halved after 30 epochs, and stopped early after 150 epochs if there was no further improvement on validation loss to prevent overfitting. Augmentation including random rotation, flipping, scaling, cropping, and brightness and contrast adjustment was applied during training.

We used the ensemble mean confidence from five deep learning models to account for variation in model predictions resulting from differences in training data among splits. We accomplished this by fine-tuning the base model with different 80/20 train/validation splits, with each split having a similar dead tree density distribution. This sampling was performed by randomly generating a list of potential splits and ranking the top five with the most similar distributions using Wasserstein distance^88^. For each training split, the model with the lowest validation loss was selected as the final model. We then used the five selected models as an ensemble, and the final detection confidence was calculated as their mean confidence. Following ref. ^88^, dead trees with mean confidence ≥ 0.35 were treated as candidate detections. The training and inference were conducted on a supercomputer using a NVIDIA A100 GPU.

Overall, the ensemble was accurate and generalized across years and regions (Extended Data Fig. 2). We evaluated the ensemble against 9,593 manually delineated dead trees in 453 plots spanning different image quality, acquisition year, mortality pattern, elevation, and land cover. At the plot level (counting the number of dead trees per ha), the ensemble achieved R² = 0.93 and slope = 0.98 on growing-season imagery (n = 397) and a slight drop in performance R² = 0.91 and slope = 0.95, when evaluated against all plots (n = 453).

We then evaluated per-tree localization accuracy for 8,263 dead tree labels from the growing-season imagery. The ensemble achieved precision, recall, and F1 scores of 0.76 each using a 6 m search radius to match predicted and annotated trees, indicating no strong bias toward commission or omission errors. However, accuracy was reduced for trees that are highly-stressed or in early-stage mortality and for trees with faint to no visible shadows owing to unfavorable sun–sensor geometry or low canopy height, which are conditions that similarly challenge human labelers (Supplementary Fig. 4). Using 1,000 bootstrap replicates, we estimated an overall model bias of −1.4% (95% CI: −8.9 to 5.7%) when evaluated against all labelled dead trees.

### Tracking individual tree mortality continuously for a decade

To create a wall-to-wall map of tree mortality rate for our study area, we tracked every detected dead tree across all observation years (Extended Data Fig. 3), and computed the total number of newly detected dead trees per year at 1-ha scale. We then overlaid the mortality rate product on the 2010 NLCD landcover map to quantify mortality across wetland and non-wetland forests. Since NAIP acquisition timing and quality vary across states and years, we developed a systematic approach to filter low-confidence mortality detections, first at the individual tree level using the multi-year record, then at the pixel level to account for varying image quality across the study domain. The detailed mortality-tracking workflow is described herein.

For each detected dead tree, we evaluated its detection timing relative to the local growing season and assigned a quality rating (hereafter acquisition quality) based on this timing. We determined the growing season from the 500 m Moderate Resolution Imaging Spectroradiometer (MODIS) phenology product, filling sparsely forested areas with the mean phenology of forested pixels at 0.5° resolution. A dead tree detected between mid-green-up and mid-green-down^91^, or if it is likely an evergreen species (according to MODIS forest type^92^ or mangrove maps^93^ in 2010) was rated as high acquisition quality. In addition, detections intersecting with defoliation polygons drawn from aerial detection surveys were ignored^78^. These criteria reduce the likelihood of misclassifying seasonally senescent deciduous or temporarily defoliated trees as dead.

We then sequentially matched detected dead trees across years. We considered a detection the same tree if a corresponding detection occurred within 6 m in any subsequent year, which is the positional uncertainty of NAIP. Each matched tree’s position was updated to the midpoint between the two detections, and its detection confidence, year, and acquisition quality were retained. In contrast, detections with no match were classified as new mortality. We leveraged parallel computing to enhance the processing speed of tracing hundreds of millions of mortality detections. Since detection confidence varied year-to-year for individual trees, we aggregated per-year probabilities into a single overall confidence score (*P_overall_*):

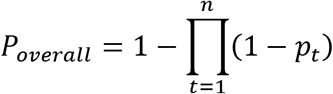

Where *p_t_* is the detection confidence in the year *t*, and *p_t_* = 0 for low acquisition quality detections. Only dead trees with *P_overall_* ≥ 0.5 were kept as valid dead trees for subsequent analyses. This threshold was selected since it aligns with forest inventory-based mortality rate estimates in regional scale (Extended Data Fig. 5). Furthermore, sensitivity analysis confirmed that the *P_overall_* threshold mostly affects the total and mean mortality rates but has little effect on the spatial pattern or temporal trend of mortality (Supplementary Fig. 5, 6), indicating the robustness of the approach. This framework enables us to confidently identify new mortality in each observation year. See ‘Sources of uncertainty’ section for factors that impact the detection.

Occasional low-quality NAIP imagery can suppress overall per-tree detection confidence across a tile (Extended Data Fig. 3), potentially leading to underestimation of mortality attributable to a given year. To account for this variation in image quality, we calculated the 75^th^ percentile of per-tree detection confidence within each 1 km grid cell for each year, and used this value as a cell-level quality score. To establish a threshold for identifying low-quality cells, we independently computed the same 75^th^ percentile confidence score for 49,674 dead trees across 10 low-quality NAIP tiles identified by human inspectors. The resulting mean ± s.d. was 0.51 ± 0.04. We therefore flagged cells with quality scores ≤ 0.5 as low-quality, resulting in approximately 32 ± 11% of cells being classified as low-quality annually.

Rather than discarding mortality detections from low-quality cell-years, we retained these mortality counts but reassigned them to the subsequent high-quality cell-year. The low-quality cell-year was treated as ‘missing’, such that the mortality count was retained, but its exact timing was not assigned to that year. This approach preserves the mortality while acknowledging that its exact timing cannot be reliably determined from the low-quality imagery. The resulting valid mortality detections were aggregated to 100 m resolution maps of new mortality per observation year.

To derive annual mortality rates from these maps, we assumed that mortality rates remained constant during years with ‘missing’ observations (resulting from absent or low-quality imagery). Observed mortality counts were then distributed evenly backward across all time steps since the previous valid observation.

Finally, for subsequent analyses, we (1) masked burned areas per year using LANDFIRE^94^ to exclude fire-induced mortality; (2) excluded the years 2010 and 2011 since they are the baseline and it is impossible to determine mortality rate without pre-baseline data; and (3) retained only pixels with a mean detection frequency greater than one year, ensuring that pixels had repeated observations on average and reducing the likelihood of false-positive mortality detections. This approach produced wall-to-wall annual maps of tree mortality rates at 100 m resolution across the coastal US from 2012 – 2023.

### Sources of uncertainty

While our approach monitored tree mortality at unprecedented spatial and temporal scale, several caveats remain. First, NAIP acquisition timing varies substantially across states and years, and in some regions — notably Florida, where most imagery was acquired in December or January — growing-season coverage is limited. Although filtering deciduous forests using land cover maps would partially address this limitation, most available products do not distinguish forest types within wetlands. We therefore relied on MODIS land cover, which, to our knowledge, is the only large-scale product that provides wetland forest type information, despite its coarser spatial resolution. This approach can therefore increase the likelihood of misclassifying deciduous trees as dead in these areas, especially for inland wetlands smaller than the MODIS resolution. This potential misclassification does not change our main conclusion about inland wetlands because a large portion of inland wetlands are distributed in the Great Lakes region. Second, the biennial revisit frequency of NAIP, combined with occasional non-growing-season or low-quality acquisitions, introduces temporal uncertainty of up to several years in mortality timing. Annual mortality rate at the local scale should therefore be interpreted with caution. Furthermore, since *P_overall_* is based on multi-year observations and becomes more reliable as additional years of observations become available, recent detections may acquire lower *P_overall_* than earlier ones. Sensitivity analysis suggested that reported mortality in recent years is more sensitive to the overall confidence threshold, but the directions of the regional trends are consistent and robust (Supplementary Fig. 6).

### Analyzing drivers of tree mortality hotspots

To examine factors that explain the formation of mortality hotspots, we employed a random forest binary classification model to predict the probability of hotspot occurrence across low-lying forests on the Great Lakes and ocean coasts, as well as inland wetlands. We defined hotspots as areas with ≥ 5 dead trees ha^−1^ yr^−1^, equivalent to ≥ 60 dead trees ha⁻¹ cumulatively over the 12-year study period, approximating the 90^th^ percentile of our mortality rate product (Extended Data Fig. 4). We selected candidate geophysical, climatic, and biotic explanatory variables based on existing literature (full list of variables can be found in Supplementary Table 2). To minimize the impacts of collinear predictors on driver attribution, predictor pairs exhibiting Pearson’s correlations (R > 0.7) were identified, and only one of each pair was retained^95^. The final predictors vary for the three flood-prone systems and with variance inflation factors (VIF) < 5, but generally include: elevation, slope, topographic position index, topographic wetness index, soil texture, soil hydric fraction, cation exchange capacity, distance to drainage, drainage density, distance to coast, salinity of the nearest coastal water body, windspeed, drought index, background climate (precipitation), extreme climate (extreme precipitation and temperature), forest type, and tree cover (Supplementary Table 3). All temporal variables were derived from the highest-resolution dataset we could find between 2012 and 2023. All layers were resampled to 100 m to align with the mortality product before model training. To mitigate spatial autocorrelation and improve computational efficiency, we extracted 25 bootstrapped subsets for the low-lying and high-elevation systems, respectively. Each low-lying and high-elevation bootstrapped subset contains 42,347 and 64,726 samples, with global Moran’s I of 0.10 (± 0.002 s.d.) and 0.07 (± 0.001 s.d.), respectively, indicating weak spatial autocorrelation at the regional scale. We then trained 25 models with the bootstrapped samples using an 80/20 train/test split and fivefold spatial cross-validation. The sizes of the spatial blocks for each system were set based on the range at which spatial autocorrelation is negligible, as identified with semi-variograms. To account for class imbalance, we assigned the minority hotspot class a higher weight based on its ratio to the non-hotspot class. Hyperparameters (number of random variables to split node and minimum node size) were optimized to maximize the F1 score. The model with the highest F1 score was selected to compute evaluation metrics. We measure model performance using Area Under the Receiver Operating Characteristic Curve (AUROC, 0-1) and Area Under the Precision-Recall Curve (AUPRC, 0-1), which measure the model’s classification ability across all probability thresholds. Higher AUROC and AUPRC values indicate better classification performance. Next, we computed variable contributions using permutation importance, which measures the drop in model accuracy when predictors are shuffled^96^. We then assessed marginal variable contributions with partial dependence plot^97^. Uncertainty in variable importance and partial dependence was measured using 95% CIs (mean ± 1.96 × s.d. of means) across the 25 models trained on bootstrapped subsets. The ‘blockCV’, ‘caret’ and ‘pdp’ packages in R (4.5.2) were used.

### Comparing high-elevation wetlands and terrestrial forests

To robustly compare inland wetlands and non-wetlands, we applied propensity score matching to locate forested pixels with similar characteristics, such as tree cover and density, climate, and topography, to minimize selection biases. To minimize spatial autocorrelation, we used the 25 bootstrapped subsets to match wetland and non-wetland locations using the ten most significant explanatory variables from the random forest model, except for variables directly affecting soil water retention potential, including soil texture and soil hydric fraction. We conducted the matching separately for hurricane-impacted zones (cumulative windspeed > 0; Supplementary Table 2) and non-impacted zones. For each subset, propensity score matching was performed using logistic regression and a nearest-neighbor approach without replacement, generating approximately 4,500 and 14,000 paired locations for impacted and non-impacted zones, respectively. Note that impacted zones should be interpreted with caution since they are mostly from suboptimal data in Florida. We ensured the site matching performance is robust (Supplementary Fig. 7). Finally, the relationship between the matched locations and environmental variables such as extreme precipitation was analyzed. Uncertainty of the analysis was quantified with 95% CI (mean ± 1.96 × s.d. of means) of the 25 bootstrapped subsets. The ‘matchit’ package in R (4.5.2) was used for the matching.

## Acknowledgement

This work is supported by the NASA Coastal Resilience Team (80NSSC23K0127). We would also like to thank Thomas S. Lever, Mahin Ganesan, Nicolas R. Miller, and Brendan D. Jalali from the School of Data Science at the University of Virginia (UVA) for their help in the initial training site selection and algorithm conceptualization. We thank the UVA Research Computing for providing the computational resources that made this work possible, as well as the Fairfax Marine Fund Endowment for providing a workstation for data analysis.

## Author contributions

H.C.H.Y. and X.Y. designed the study with highly valuable input from all co-authors. H.C.H.Y. performed all the analyses. H.C.H.Y. and X.Y. prepared the initial manuscript, with advice and discussions from all authors. All authors contributed to the writing of the paper.

## Extended Data

**Extended Data Table 1.**
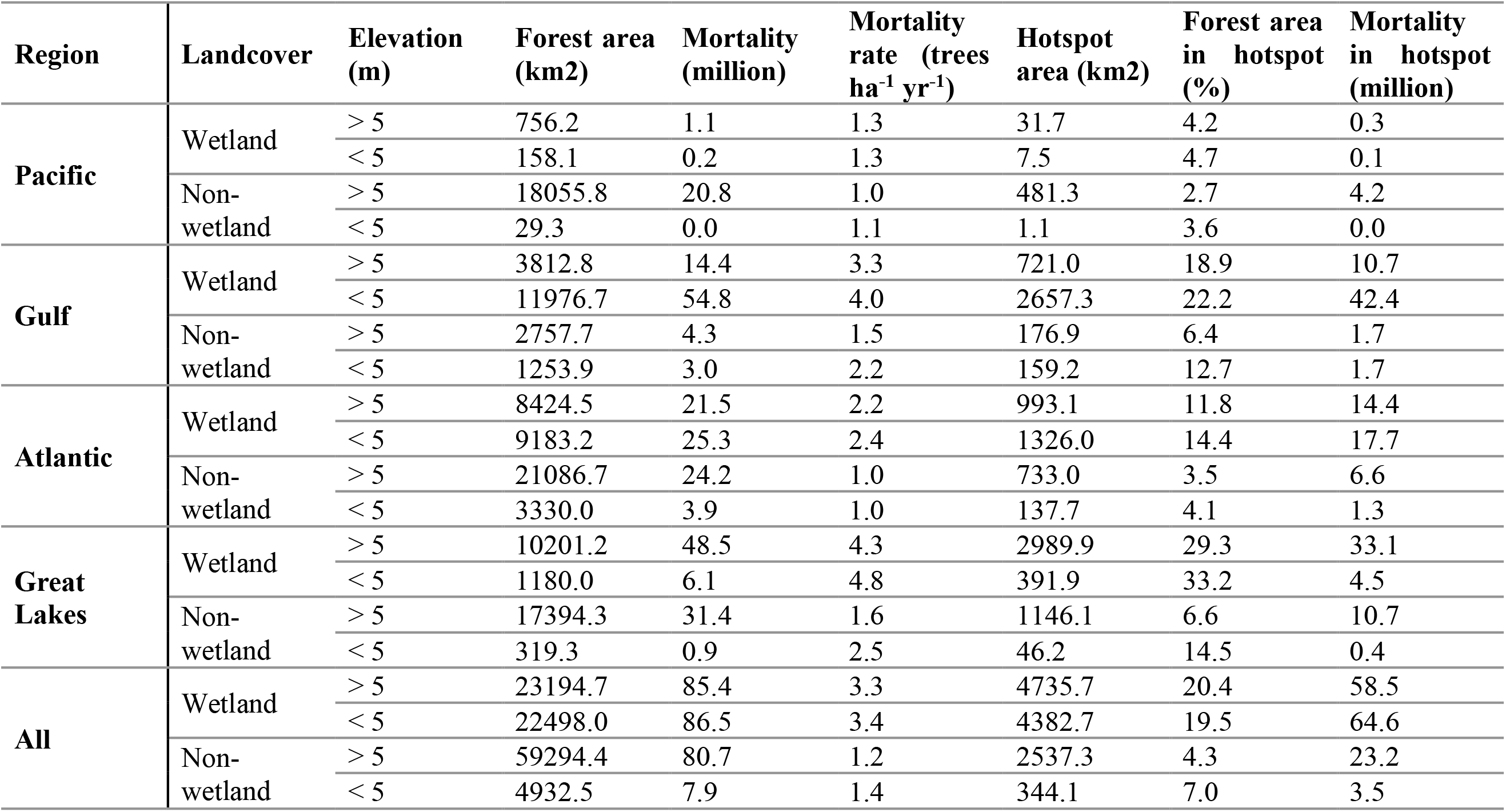
Tree mortality and hotspot distribution across the US coastal forests mapped between 2012 and 2023.

| Region | Landcover | Elevation<br>(m) | Forest area<br>(km2) | Mortality<br>(million) | Mortality<br>rate (trees<br>ha<sup>-1</sup> yr<sup>-1</sup>) | Hotspot<br>area (km2) | Forest area<br>in hotspot<br>(%) | Mortality<br>in hotspot<br>(million) |
| --- | --- | --- | --- | --- | --- | --- | --- | --- |
| Pacific | Wetland | > 5 | 756.2 | 1.1 | 1.3 | 31.7 | 4.2 | 0.3 |
|  |  | < 5 | 158.1 | 0.2 | 1.3 | 7.5 | 4.7 | 0.1 |
|  | Non-wetland | > 5 | 18055.8 | 20.8 | 1.0 | 481.3 | 2.7 | 4.2 |
|  |  | < 5 | 29.3 | 0.0 | 1.1 | 1.1 | 3.6 | 0.0 |
| Gulf | Wetland | > 5 | 3812.8 | 14.4 | 3.3 | 721.0 | 18.9 | 10.7 |
|  |  | < 5 | 11976.7 | 54.8 | 4.0 | 2657.3 | 22.2 | 42.4 |
|  | Non-wetland | > 5 | 2757.7 | 4.3 | 1.5 | 176.9 | 6.4 | 1.7 |
|  |  | < 5 | 1253.9 | 3.0 | 2.2 | 159.2 | 12.7 | 1.7 |
| Atlantic | Wetland | > 5 | 8424.5 | 21.5 | 2.2 | 993.1 | 11.8 | 14.4 |
|  |  | < 5 | 9183.2 | 25.3 | 2.4 | 1326.0 | 14.4 | 17.7 |
|  | Non-wetland | > 5 | 21086.7 | 24.2 | 1.0 | 733.0 | 3.5 | 6.6 |
|  |  | < 5 | 3330.0 | 3.9 | 1.0 | 137.7 | 4.1 | 1.3 |
| Great Lakes | Wetland | > 5 | 10201.2 | 48.5 | 4.3 | 2989.9 | 29.3 | 33.1 |
|  |  | < 5 | 1180.0 | 6.1 | 4.8 | 391.9 | 33.2 | 4.5 |
|  | Non-wetland | > 5 | 17394.3 | 31.4 | 1.6 | 1146.1 | 6.6 | 10.7 |
|  |  | < 5 | 319.3 | 0.9 | 2.5 | 46.2 | 14.5 | 0.4 |
| All | Wetland | > 5 | 23194.7 | 85.4 | 3.3 | 4735.7 | 20.4 | 58.5 |
|  |  | < 5 | 22498.0 | 86.5 | 3.4 | 4382.7 | 19.5 | 64.6 |
|  | Non-wetland | > 5 | 59294.4 | 80.7 | 1.2 | 2537.3 | 4.3 | 23.2 |
|  |  | < 5 | 4932.5 | 7.9 | 1.4 | 344.1 | 7.0 | 3.5 |

**Extended Data Fig. 1.**
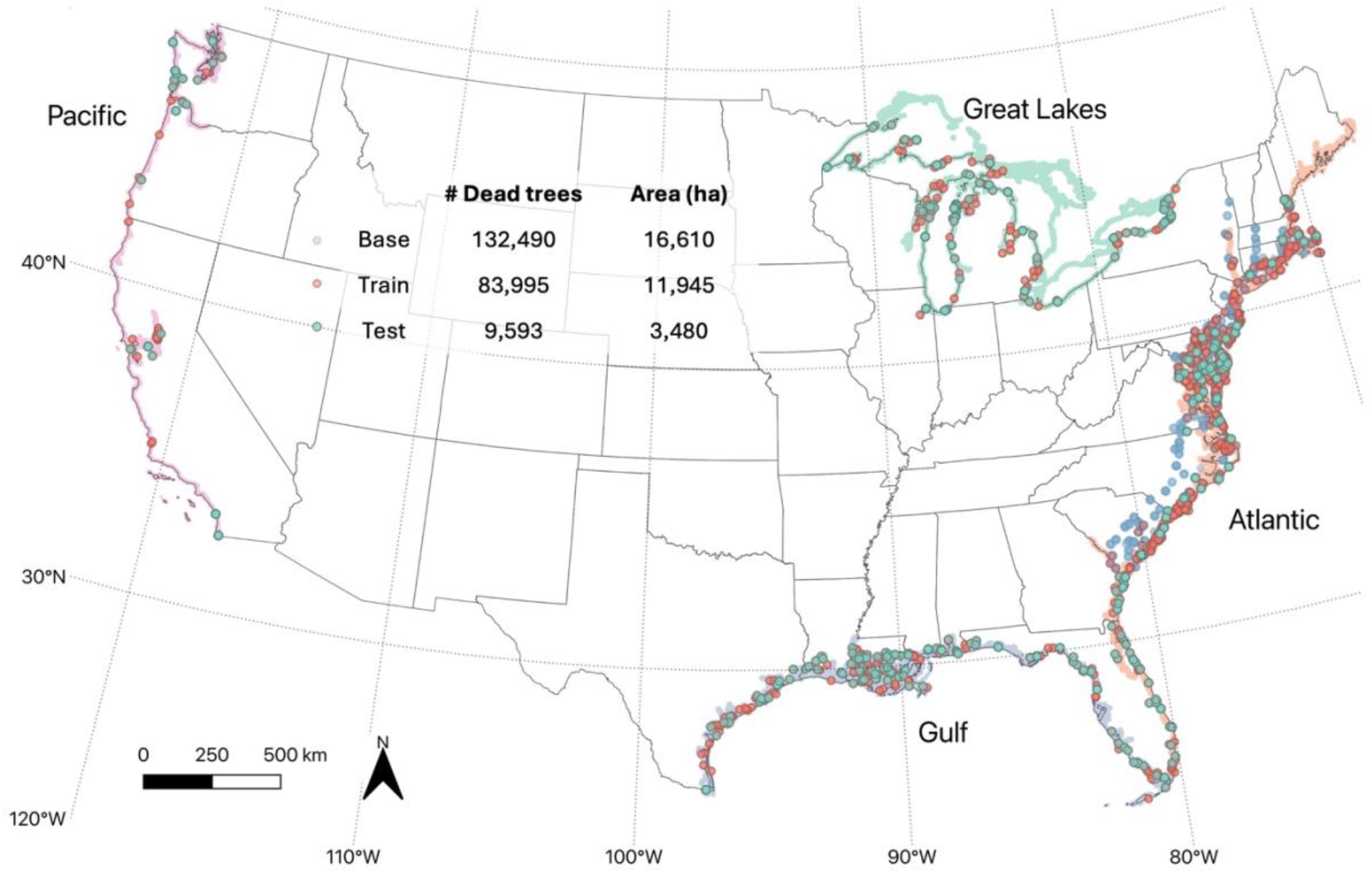
Overview of training and test sites. The locations of training and test plots used for model training. The training involves 216,485 dead tree labels (132,490 labels from the Atlantic coast for base model training; 83,995 labels from all coastlines for fine-tuning). The fine-tuning labels are collected from 2010 to 2023. The final model was tested on an independent dataset consisting of 9,593 manually labeled dead trees covering 3,480 ha. State boundaries from the Census Bureau.

**Extended Data Fig. 2.**
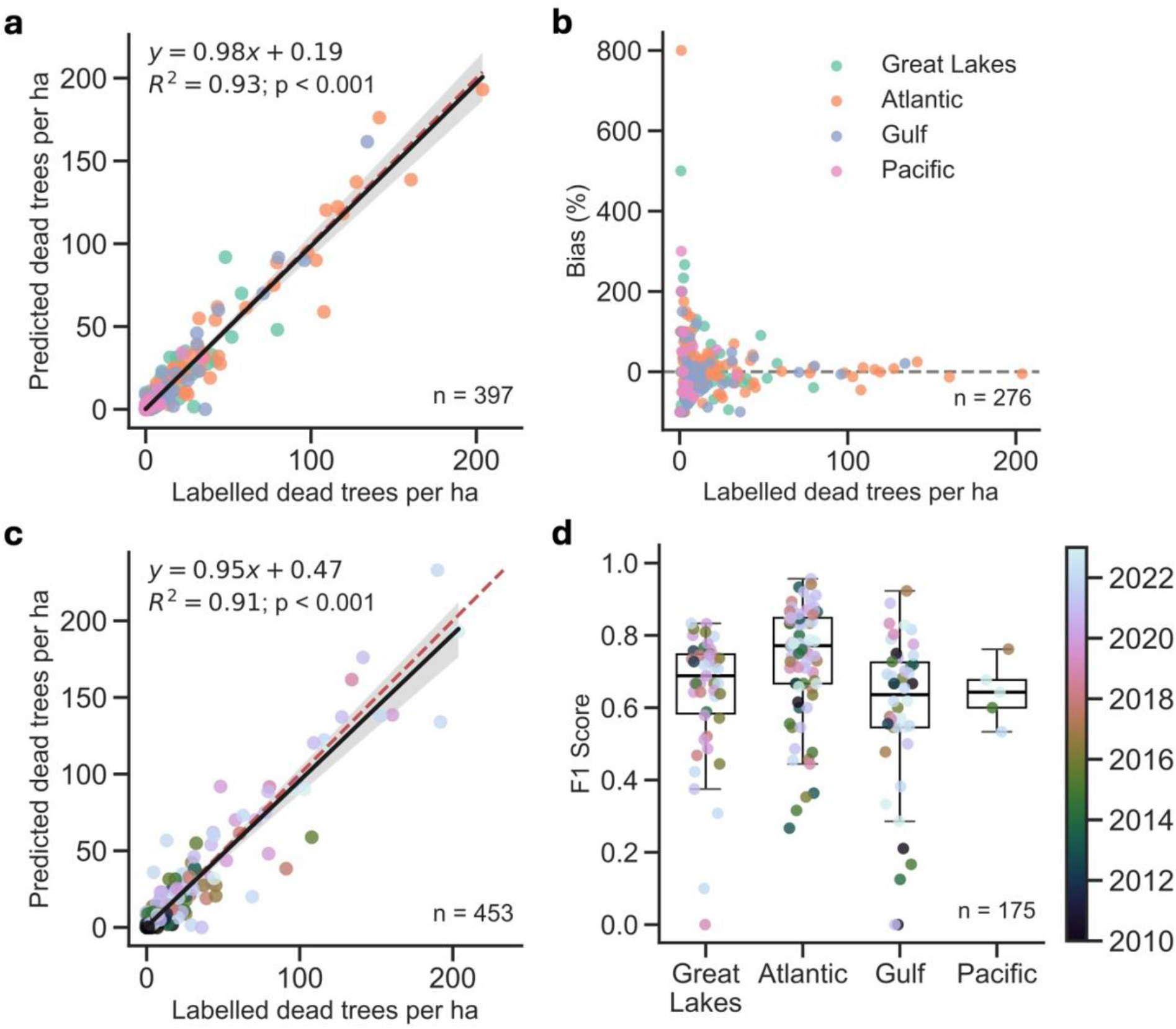
Model performance in detecting individual dead trees. We compared 9,593 manually labeled dead trees with heatmap-model predictions across 453 plots spanning all U.S. coasts from 2010 to 2023. **a**, Comparison between predicted and labeled dead trees per hectare, considering only growing season detections (n = 397) as determined by the MODIS phenology product. Colors denote different coasts, and the shaded region indicates the 95% CI. Dashed red line indicates the 1:1 line. **b**, Percent bias of individual plots, where deviations decrease with increasing dead tree density. **c**, As in **a**, but colored by image acquisition year and including all 453 plots, including those outside the growing season. **d**, Overall localization accuracy (F1 score) of individual detections with a search radius of 6 m. Each point represents the F1 score of one plot. The box shows the median (centerline) and interquartile range (bounds), and whiskers extend to 1.5 × the interquartile range. Only plots with > 5 labeled dead trees are shown, since F1 scores are highly sensitive at low sample sizes. Color denotes acquisition year. The final product only retained growing season detections. We acknowledge model performance can decline when the scene is equally difficult for human labelers to locate dead trees (Supplementary Fig. 4).

**Extended Data Fig. 3.**
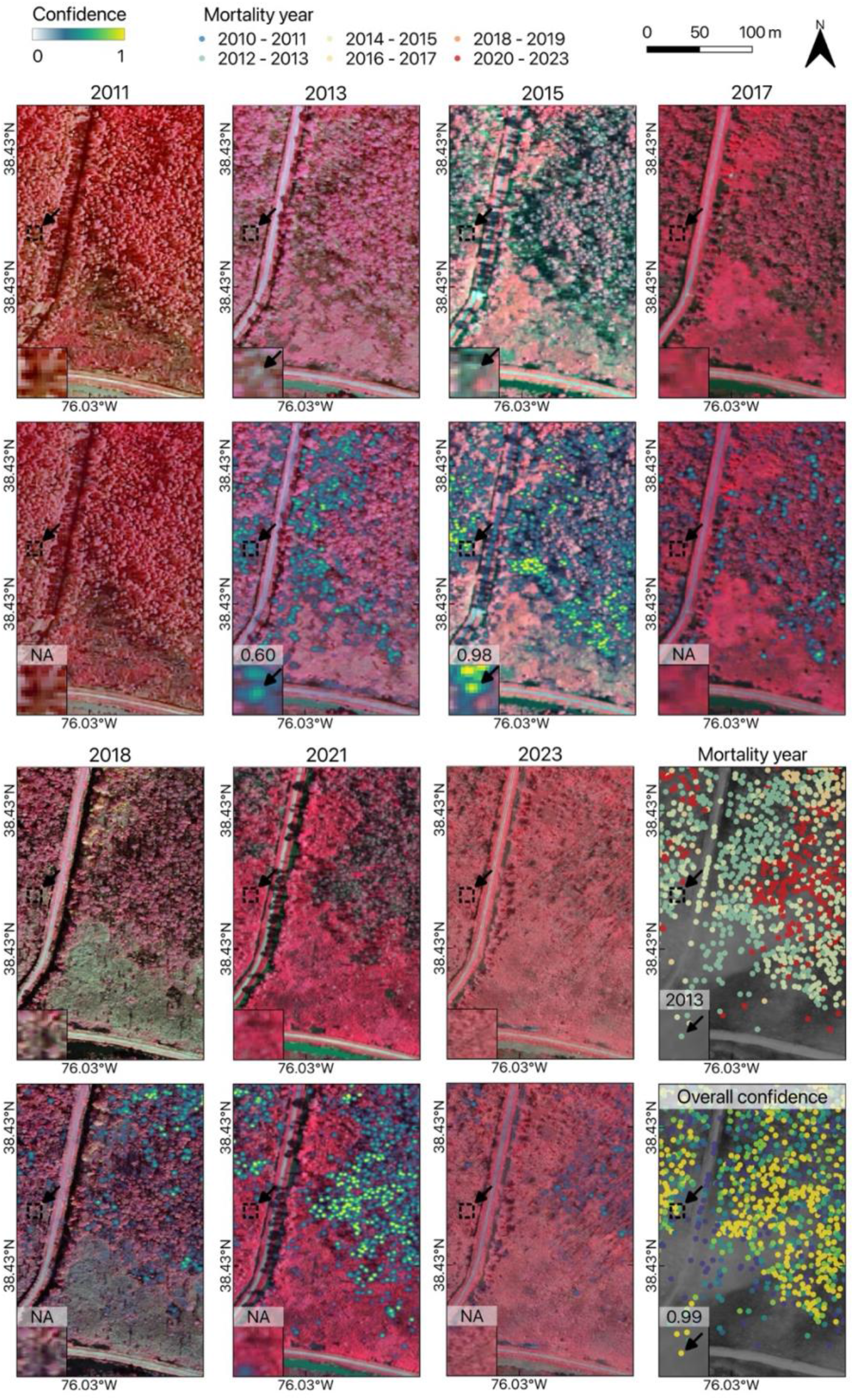
Tracking mortality timing biennially with heatmap predictions. Example of NAIP images and heatmap predictions from 2011 to 2023. The mortality year represents the year when the dead tree is first detected, while the overall confidence was computed as the cumulative confidence across all years (Methods). We only retained detections with overall confidence ≥ 0.5 for subsequent analyses. Basemaps © NAIP and ESRI.

**Extended Data Fig. 4.**
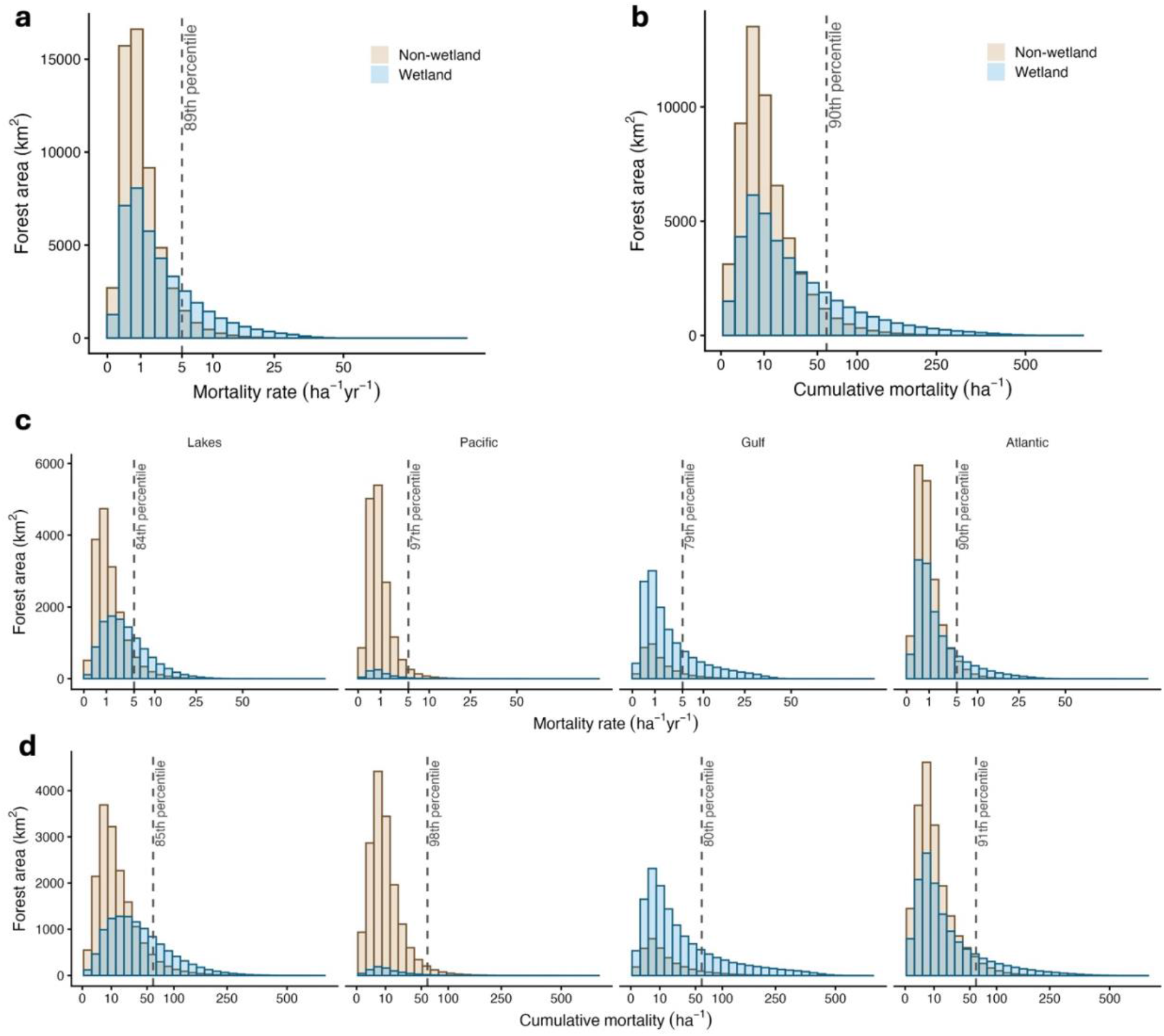
Histograms of mortality rate and cumulative mortality across the US coasts. Histograms of forest area by mortality rate (**a**, **c**; trees ha⁻¹ yr⁻¹) and cumulative mortality (**b**, **d**; trees ha⁻¹) for wetlands and non-wetlands. **a**, **b**, All regions combined. **c**, **d**, Regions shown separately. Dashed lines indicate the mortality hotspot threshold (5 trees ha⁻¹ yr⁻¹, equivalent to ∼60 trees ha⁻¹ cumulatively from 2012 to 2023). Labels show the percentile of each distribution corresponding to this threshold.

**Extended Data Fig. 5.**
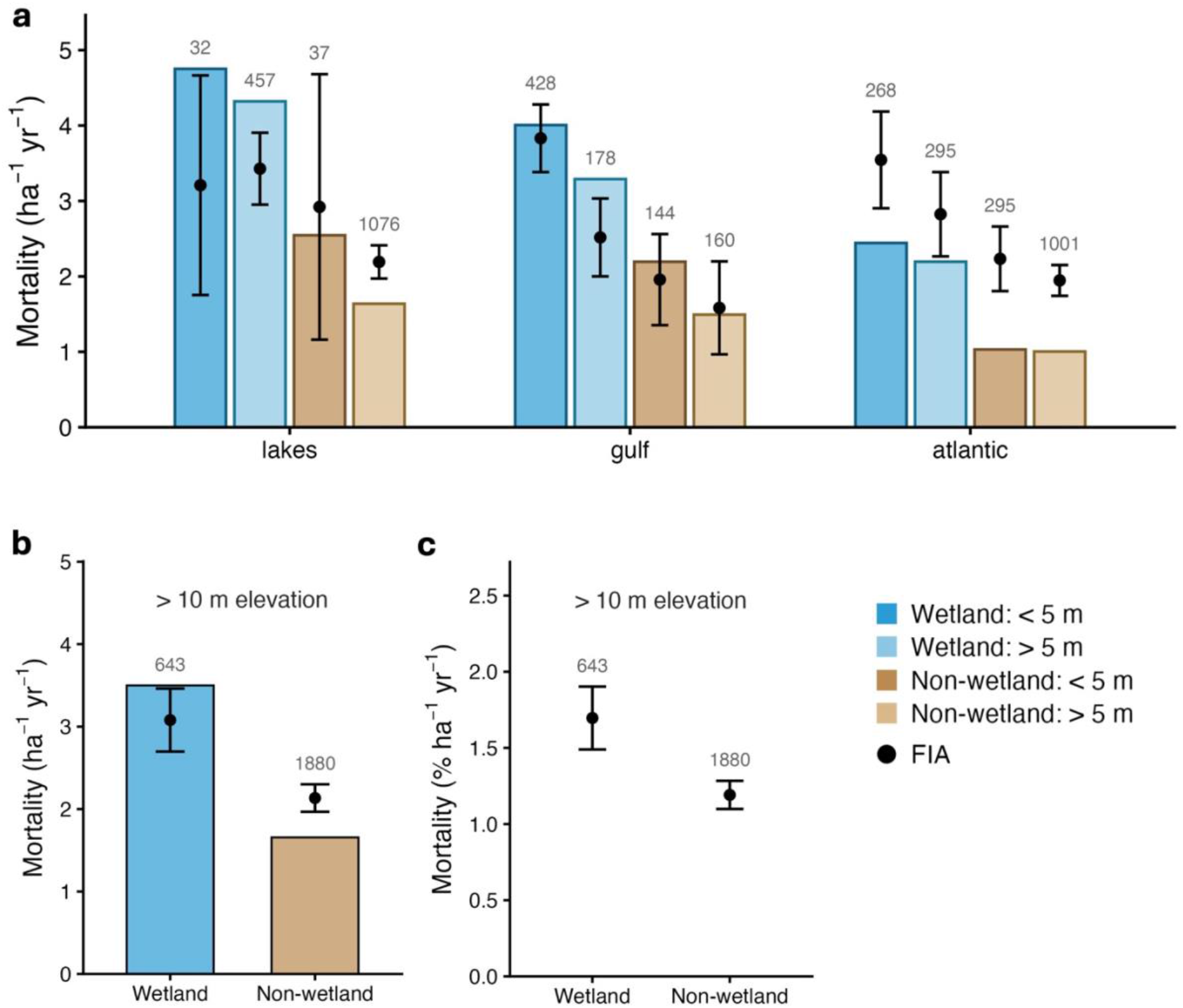
Comparison with forest inventory data. Mortality rate of canopy trees (diameter ≥ 20 cm) estimated from paired re-measurements of Forest Inventory and Analysis (FIA) plots where the most recent survey was conducted during 2012-2023, using the *growMort* function in the *rFIA* package in R. We reclassify the ‘flooded and swamp forest’ as wetland based on the National Vegetation Classification Standard (NVCS) of the plots. Note that plots from the Pacific coasts are excluded since NVCS were not assigned. **a**, Annual mortality rate across regions and elevation groups. Points and error bars indicate the mean and 95% CI of FIA estimates. Numbers above each point represent the number of FIA plots in each group. Bar heights show the mean rate calculated using our mortality product for each region. **b**, Mortality rate for high-elevation forests, and **c**, same as **b**, but accounts for live trees and represents the percent of mortality since the start of measurement periods. Since the elevations recorded by FIA are not necessarily referenced to the lake-level datum we used, we extracted the elevations of all plots from the 3DEP DEM, which can lead to potential error since the locations of the FIA plots are shuffled. However, based on the plots on the oceanic coasts, we found the difference between the 3DEP and FIA elevations is small (−0.02 ± 1.97 m s.d., n = 2,769), indicating a small effect at the regional scale.

**Extended Data Fig. 6.**
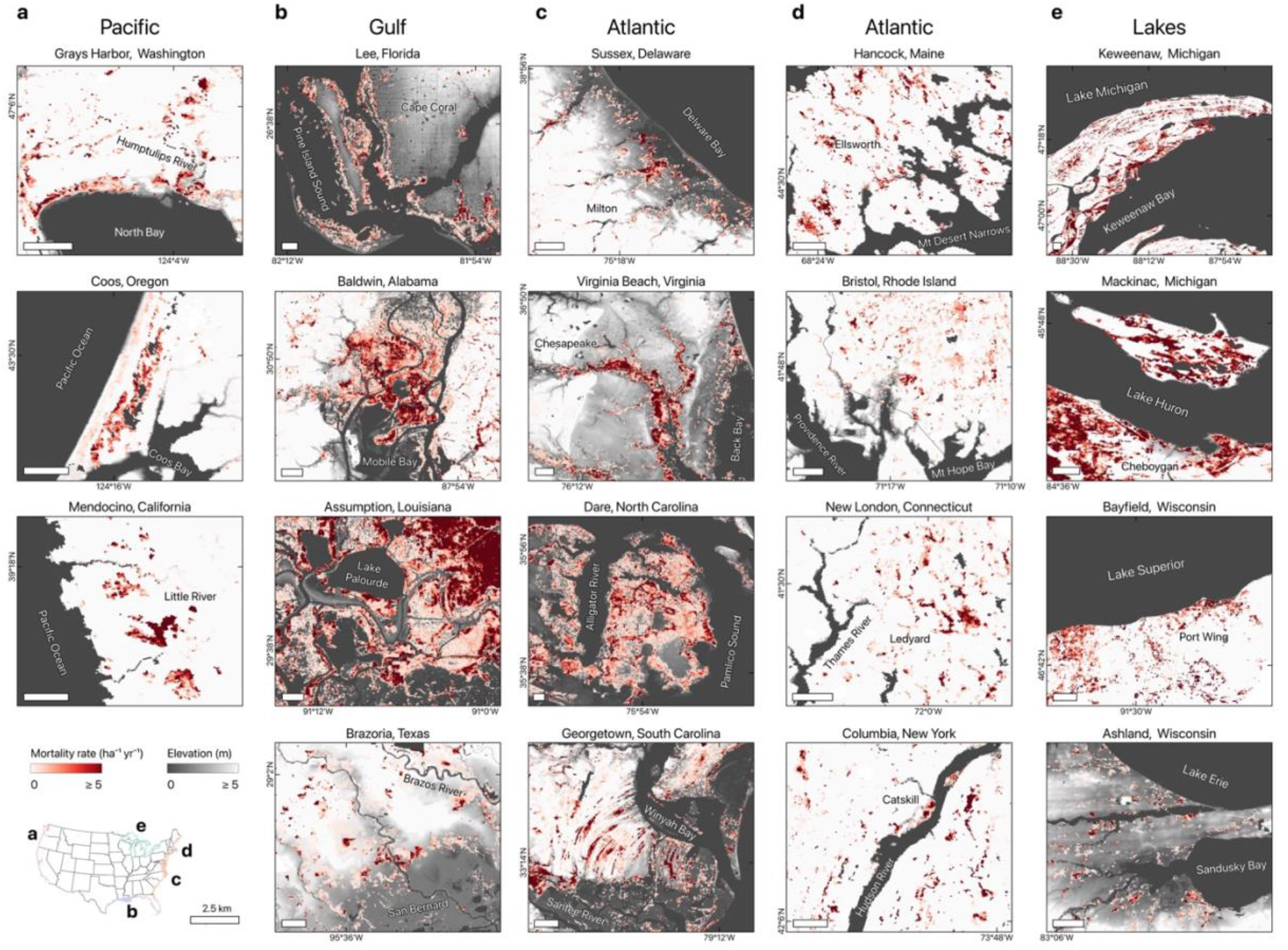
Extensive tree mortality across large and small wetland forests. Examples of tree mortality hotspots in wetland forests across the Pacific, Gulf, Atlantic coasts, and the Great Lakes. Elevations are referenced to NAVD88 for oceanic coasts and the lake-specific Low Water Datum (LWD) for Great Lakes regions, respectively. Note that not all of the mortality shown is necessarily driven by flooding or saltwater intrusion. We masked mortality associated with infestation and fire using aerial detection survey datasets, which may not cover every infestation and fire event.

**Extended Data Fig. 7.**
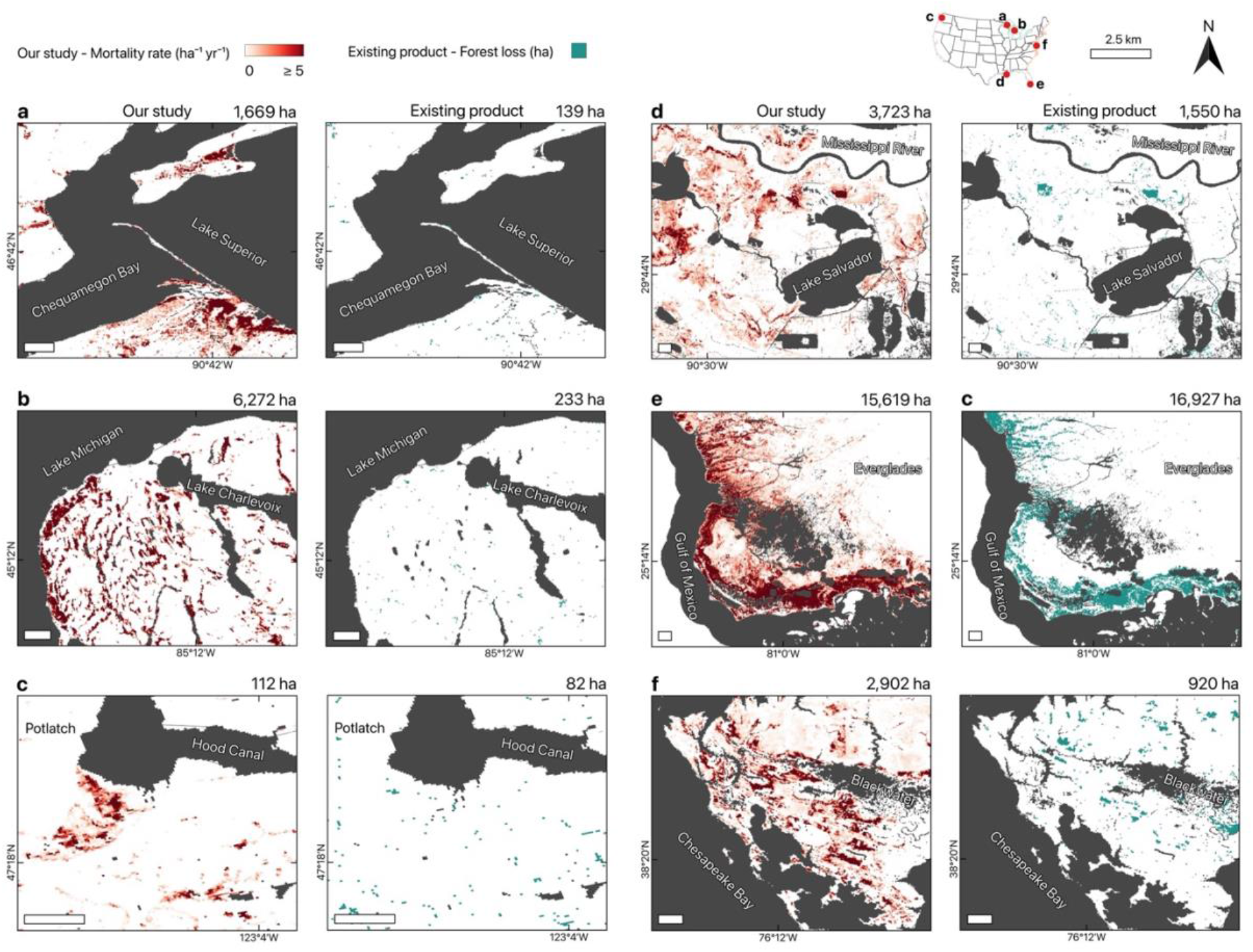
Comparison of wetland forest loss with existing products. Examples comparing our individual tree mortality map with an existing Landsat-based forest loss product from Hansen et al. We calculated all forest loss areas from 2012 to 2023, excluding human-driven land cover change (for example, agriculture and urban development) according to NLCD. Mortality and forest loss associated with infestation and fire were masked using aerial detection survey datasets. Most mortality hotspots (≥ 5 dead trees ha^−1^ yr^−1^) mapped in our study were not detected by the Landsat-based product. The values in the top-right of each panel indicate the areas of mortality hotspots detected in our product and forest loss detected in the Hansen product, respectively. Note that a direct comparison is not feasible because human-driven loss cannot be completely excluded from the Landsat-based product, particularly in plantation forests, and our approach captures a gradient of tree mortality rather than a binary forest-loss classification. While the two products measure different aspects of forest change, this comparison highlights that coarse-resolution products can substantially underestimate forest loss due to fine-scale, heterogeneous tree mortality.

**Extended Data Fig. 8.**
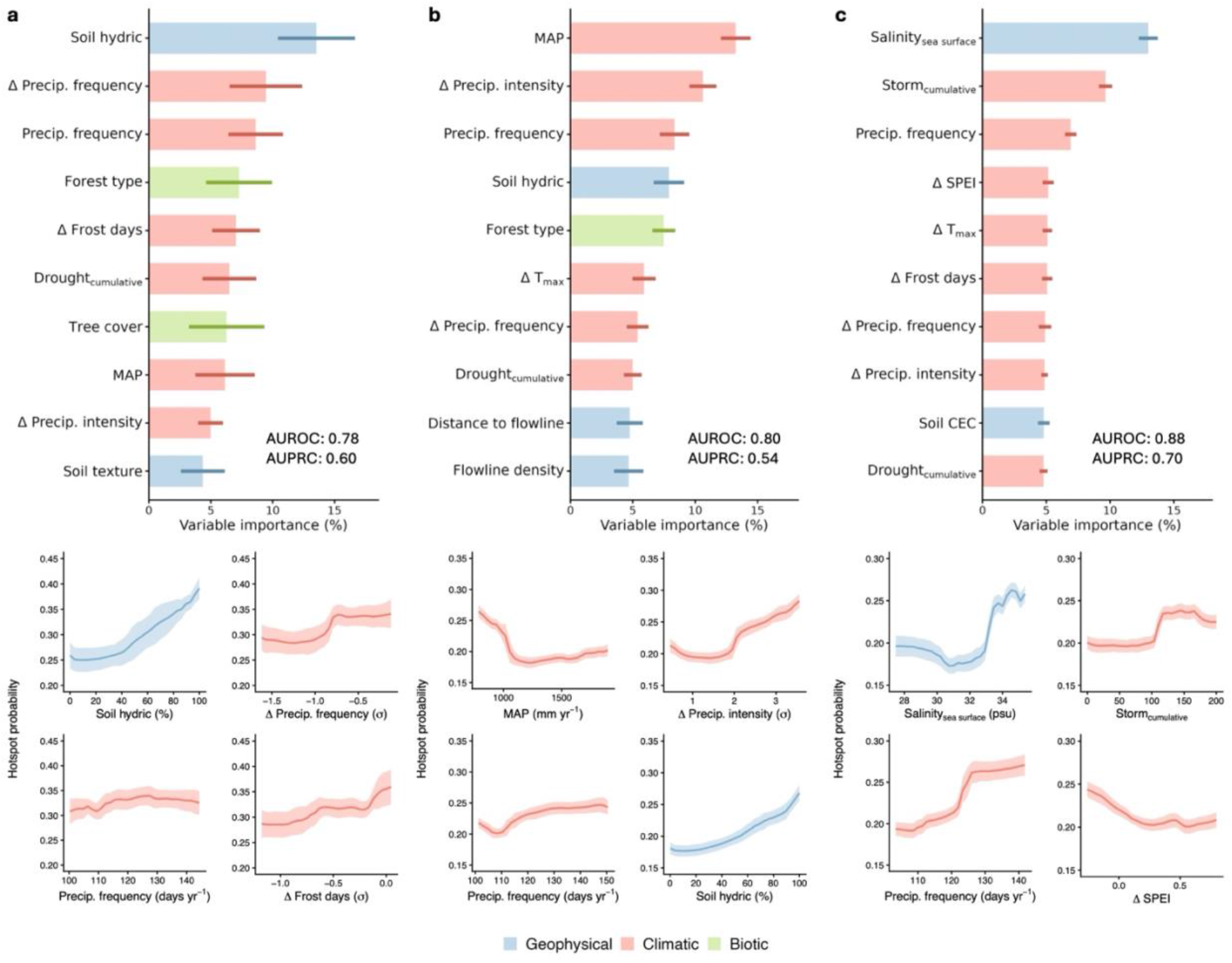
Drivers of mortality hotspot formation across flood-prone forests. Random forest models were used to explain the probability of hotspot formation across three systems (Method): **a**, Low-lying forests on lake margins (Great Lakes); **b**, Inland wetlands (> 10 m elevation; **c**, Low-lying forests on ocean coasts (Pacific, Atlantic, and Gulf). We defined hotspots as areas with ≥ 5 dead trees ha^−1^ yr^−1^ (approximately the 90^th^ percentile). Model performance is presented with AUROC and AUPRC, which range from 0 to 1. Higher AUROC and AUPRC values indicate better classification performance. The relative importance of predictor variables, model performance metrics, and partial dependence plots of the top 4 most important numeric variables in explaining mortality hotspot occurrence are presented. Bar height and Error bars in the variable importance plots represent the mean importance and the 95% CI across the bootstrapped models. The same applies to the solid line and shaded regions in the partial dependence plots. Abbreviations: cation exchange capacity (CEC), Standardized Precipitation-Evapotranspiration Index (SPEI), Annual maximum temperature (T_max_), Mean annual precipitation (MAP), Precipitation (Precip.). Details of individual variables used for each model are shown in Supplementary Table 2 and 3.

**Extended Data Fig. 9.**
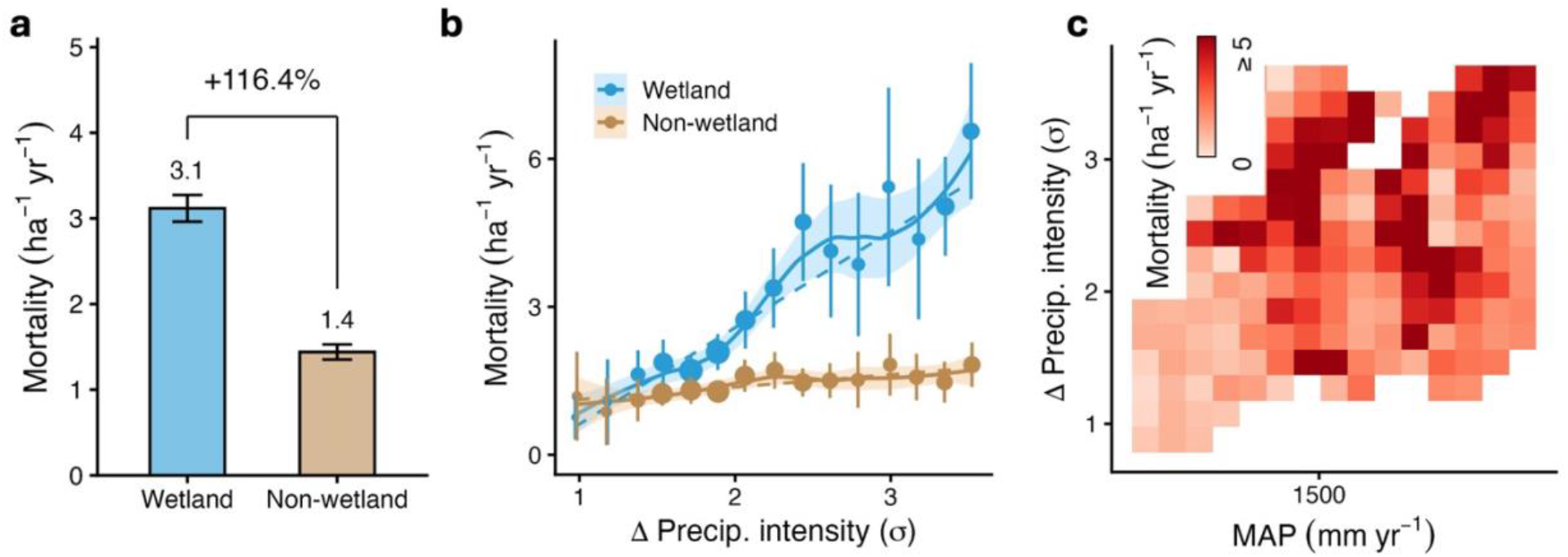
Extreme precipitation exacerbates mortality in storm-impacted inland wetlands. **a**, Comparison of wetland and non-wetland tree mortality rates after matching sites with similar characteristics within hurricane-impacted zones. The percentage indicates the relative increase in wetland mortality compared to non-wetlands. Bar height and error bar represent the mean and 95% CI of the bootstrapped subsets (n = 25; Methods). **b**, Response of wetland and non-wetland mortality rate against precipitation intensity anomaly (bin averages ± 95% CI). Point size reflects the area of forest sampled (1,000 to 25,000 ha). Dashed line is the linear regression weighted by standard error (Wetland: slope = 1.95, R^2^ = 0.91, P ≤ 0.0001; Non-wetland: slope = 0.25, R^2^ = 0.61, P = 0.0005). Solid line represents a local polynomial fit for visualization, shaded with 95% CI. **c**, Wetland mortality rate against mean annual precipitation (MAP; horizontal axis) and precipitation intensity anomaly (vertical axis). Color shows the mean mortality rate of the bootstrapped samples. In **b** and **c**, the outer 2.5% data of each tail were excluded from the distribution to ensure adequate samples in each bin. Note that many storm-impacted inland wetlands within our domain are located in Florida, where NAIP imagery is frequently acquired during December–January and is therefore of suboptimal quality (Supplementary Fig. 3). This limits our confidence in distinguishing deciduous trees in wetlands from dead, and thus should be interpreted with caution.

