## Supplementary Fig 1-7, Supplementary Table 1-3 for "Rising water is an underappreciated driver of forest mortality"

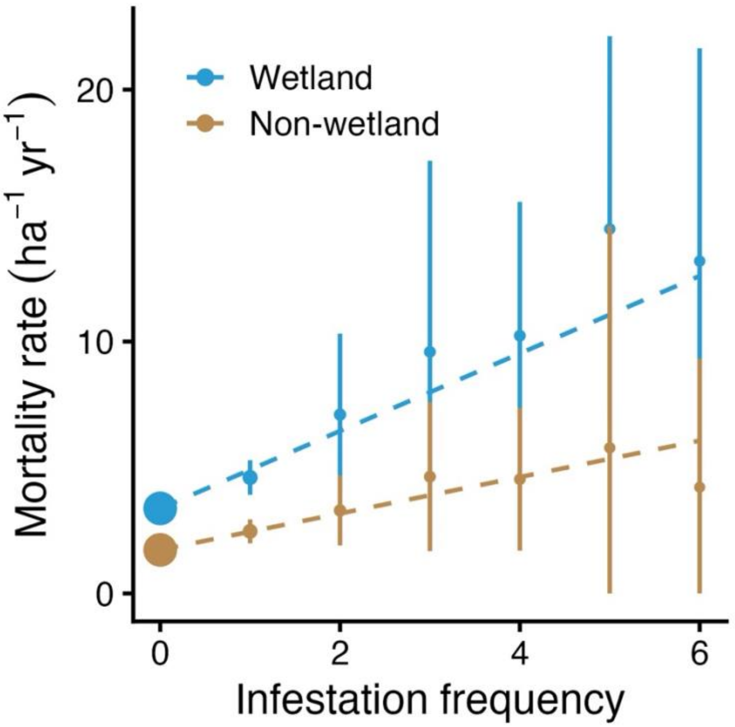


**Supplementary Fig. 1 | Tree mortality in wetlands shows a stronger relationship with insect infestation.** We compared wetland and non-wetland tree mortality rates against infestation frequency (insect and pathogens) after matching sites with similar characteristics, excluding hurricane-impacted zones (Methods). The frequency refers to the number of infestation events that occurred between 2012 and 2023. Dashed line is the linear regression weighted by standard error Point and error bar represent the mean and 95% CI of bootstrapped subsets (n = 25).

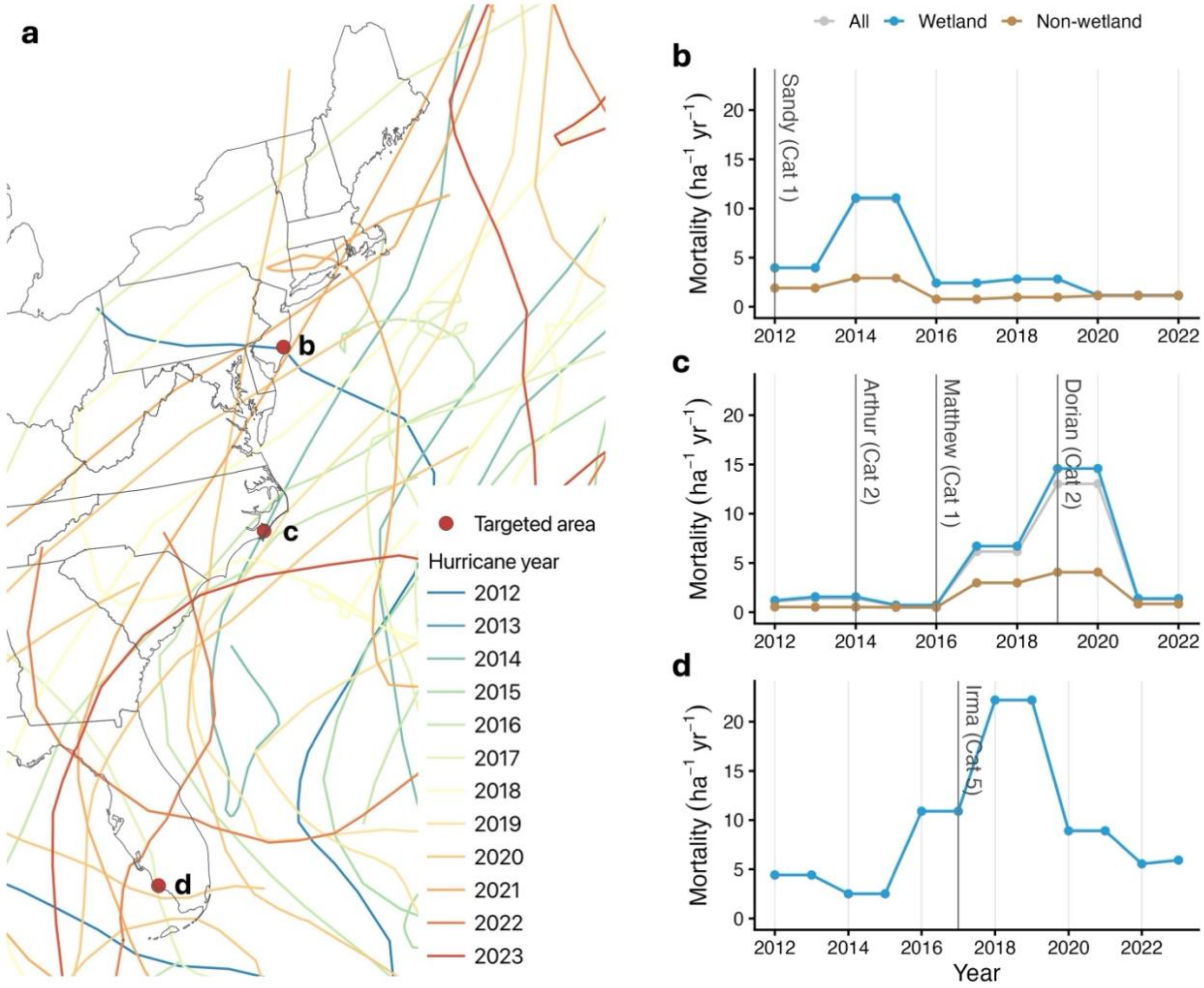


**Supplementary Fig. 2 | Localized temporal trend of tree mortality in selected areas impacted by hurricanes. a**, Map of hurricane tracks in the eastern US during 2012 – 2023. Points represent the selected locations for temporal trend analysis. **b** – **d**, We calculated average annual mortality rates of the selected locations (15 km radius buffer). The name, year, and category of hurricanes that are within 110 km (60 nautical miles) of the selected locations were labeled. **b**, Temperate forest in New Jersey, Atlantic. **c**, Temperate forest in North Carolina, Atlantic. **d**, Mangrove in Florida, Gulf. Non-wetland forests in **c** were not plotted since they occupy less than 20 ha of the area. Hurricane information was retrieved from NOAA. State boundaries from the Census Bureau.

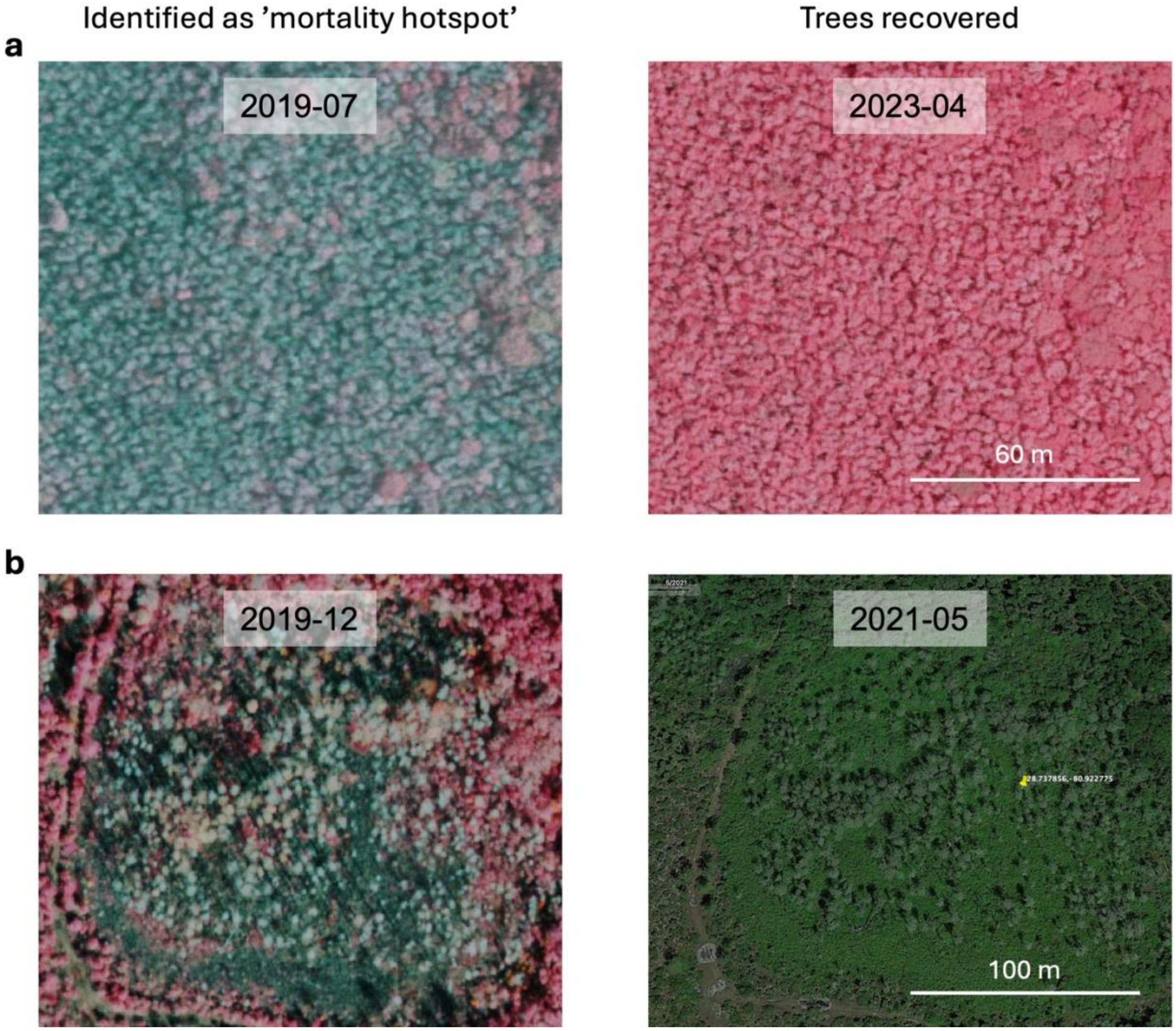


**Supplementary Fig. 3 | Recovery of wetland forests.** Examples of wetland forests showing signs of recovery, potentially related to the nonstructural carbohydrate reserves in trees to buffer them against highly stressed conditions, although interpretation is limited by image quality. **a**, low-lying hardwood forest in South Carolina. **b**, inland wetland forest recovery after a storm landfall in Florida. However, many storm-impacted inland wetlands within our domain are located in Florida, where NAIP imagery is frequently acquired during December–January and is therefore of suboptimal quality. This limits our confidence in distinguishing deciduous trees in wetlands from dead. Images from NAIP and Google Earth Pro images.

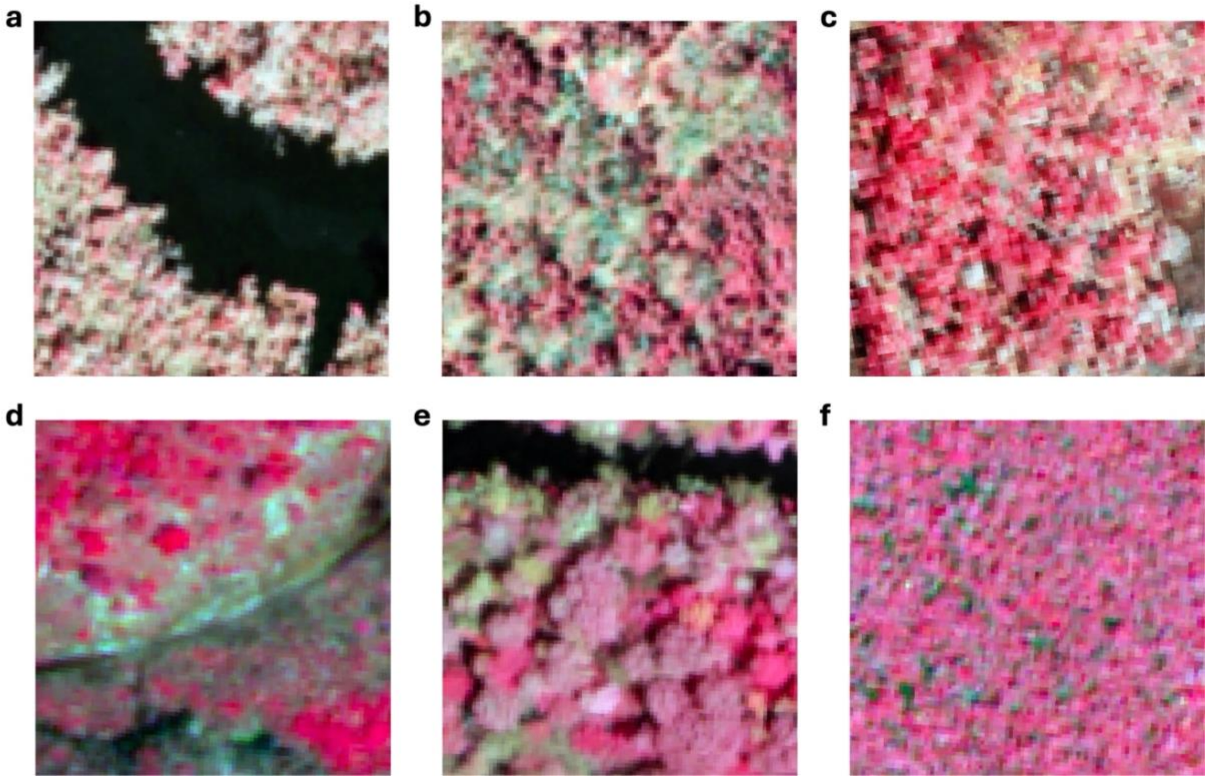


**Supplementary Fig. 4 | Examples of challenging scenarios for both models and human annotators.** Highly stressed trees, exhibiting a brown color, and those with faint or no visible shadows, often due to unfavorable sun–sensor geometry, short trees or shrubs, can pose challenges to mortality detection. Examples are from a) FL, b) MA, c) NC, d) FL, e) LA, f) FL. Images from NAIP.

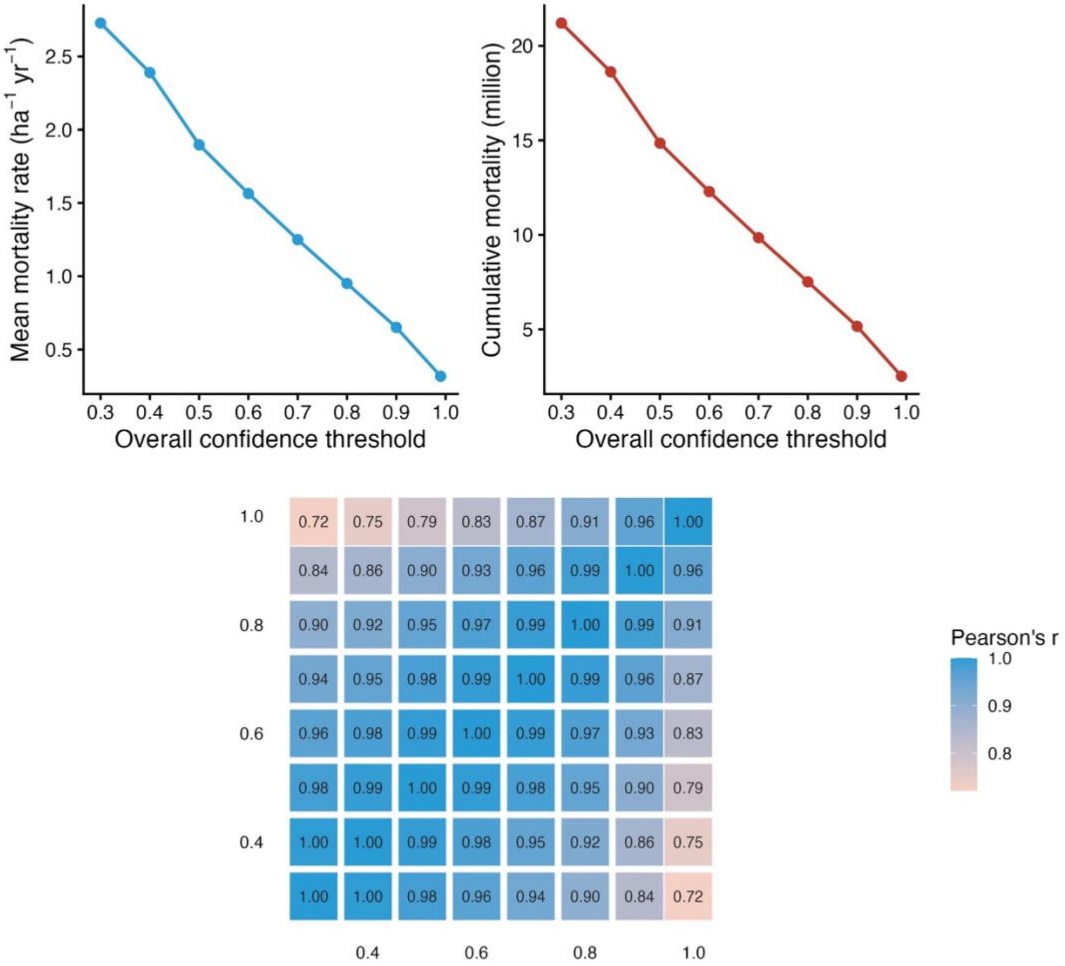


**Supplementary Fig. 5 | Sensitivity of spatial analysis to different overall confidence thresholds.** We computed the mortality products using multiple overall confidence thresholds (0.3 – 0.99), and extracted 1 million pixels to test the sensitivity of **a**, the mean mortality rate, and **b**, cumulative mortality against different thresholds. As expected, both decline monotonically as the threshold increases. **c**, Pairwise Pearson correlation of the mortality rate products between threshold pairs. High r reflects increased similarity in the spatial patterns among the pairs. This implies the spatial patterns of mortality are highly similar within the threshold range of 0.3 to 0.8, reflecting the robustness of the approach. Moreover, a final threshold of 0.5 was selected because its mean mortality rate aligned with stand-level inventory from FIA at the regional scale (Extended Data Fig. 4).

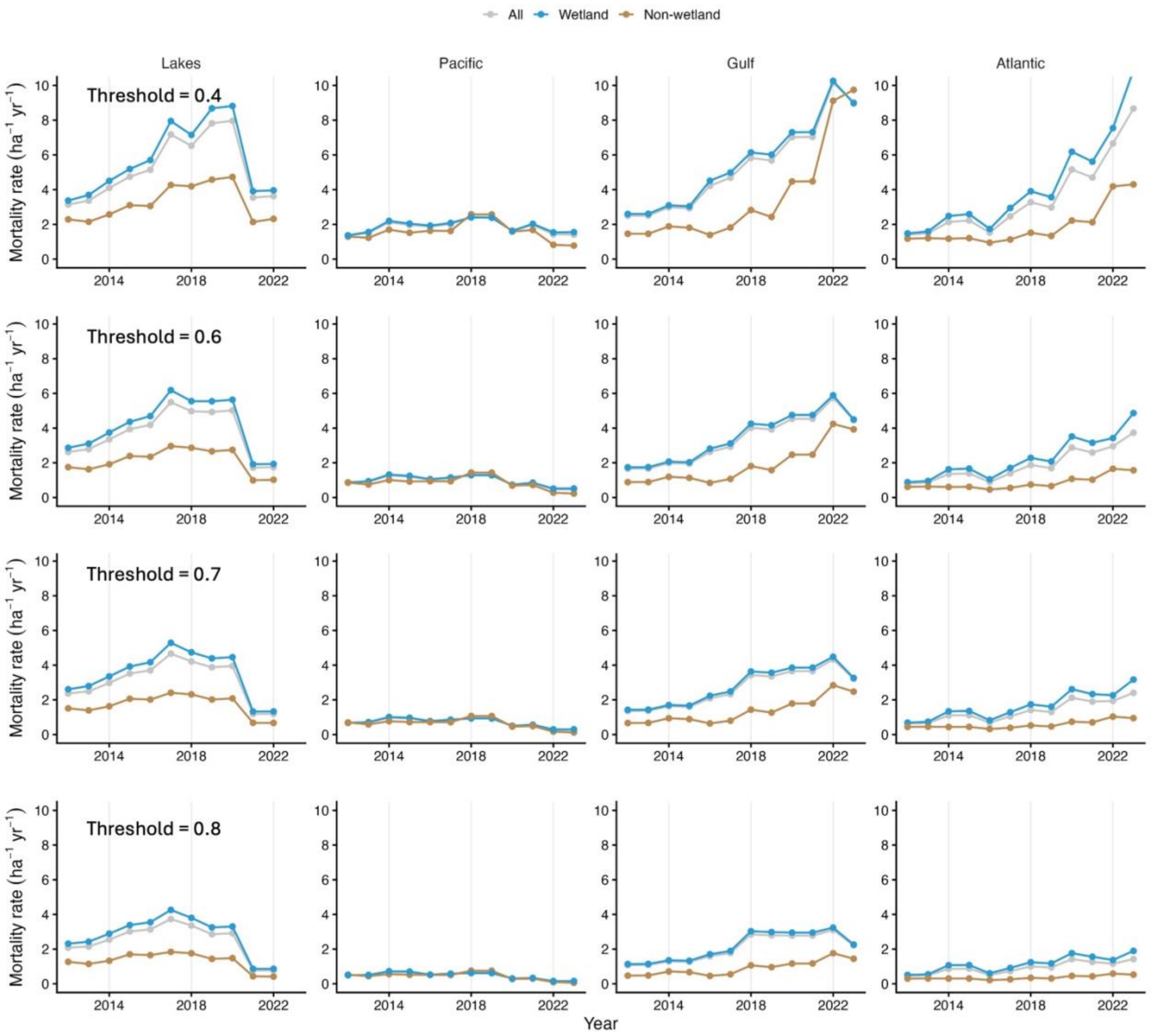


**Supplementary Fig. 6 | Sensitivity of temporal analysis to different overall confidence thresholds.** We computed the mortality products using multiple overall confidence thresholds (0.4, 0.6, 0.7 and 0.8) to test the sensitivity of the temporal trend of regional low-lying forest mortality rate against different thresholds. The decreasing slope with increasing thresholds is expected since new mortality that occurred in more recent years is less likely to have repeated detections, and hence fewer detections can satisfy the overall confidence thresholds. Nevertheless, the results indicate that the directions of the regional trends are consistent and robust across varying overall confidence thresholds.

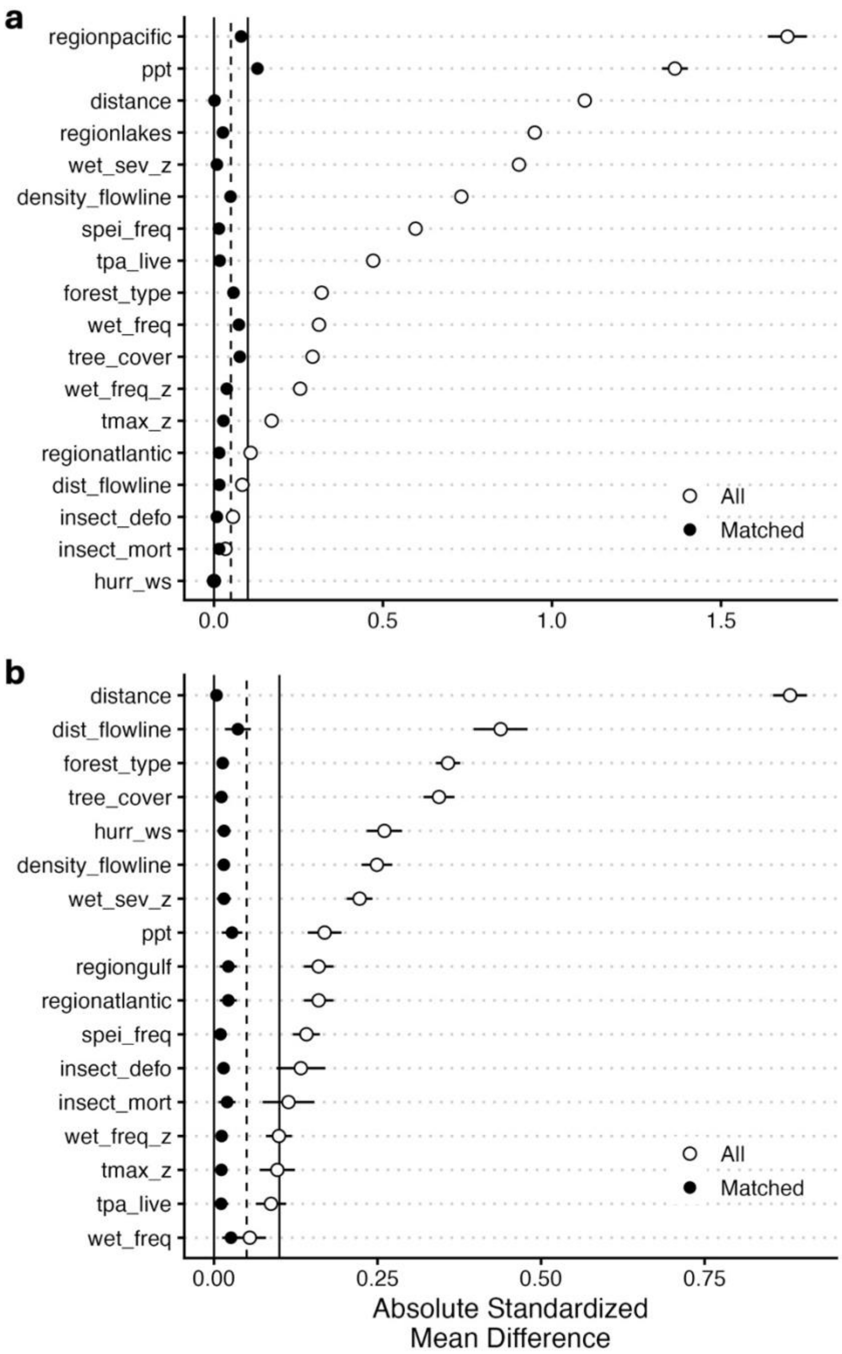


**Supplementary Fig. 7 | Site-matching performance for inland wetland and non-wetland forests**. **a**, non-storm-impacted-zone only. **b**, storm-impacted zone only. Absolute standardized mean difference (ASMD) refers to the difference between the matched and unmatched (all) data. ASMD values less than 0.1 indicate good matching. Point and error bar represent the mean and 95% CI of bootstrapped samples (n = 25).

**Supplementary Table 1** | Pearson's correlation of regional mortality rate and environmental variables.

| Region | Variable | All | Wetland | Non-wetland |
|---|---|---|---|---|
| Pacific | Water level | 0.28 | 0.34 | 0.08 |
| | Drought only | 0.19 | 0.17 | 0.24 |
| Gulf | Water level | 0.70* | 0.73** | 0.41 |
| | Hurricane only | 0.67* | 0.70* | 0.31 |
| | Drought only | -0.29 | -0.32 | -0.02 |
| | Compound | -0.29 | 0.31 | -0.14 |
| Atlantic | Water level | 0.87*** | 0.88*** | 0.75** |
| | Hurricane only | -0.01 | 0.00 | -0.07 |
| | Drought only | -0.19 | -0.20 | -0.04 |
| | Compound | -0.09 | -0.10 | -0.08 |
| Lakes | Water level | 0.68* | 0.68* | 0.62* |
| | Drought only | -0.5 | -0.5 | -0.43 |

Water level refers to averaged yearly sea- or lake-levels across gauges. Hurricanes and droughts refer to fractional areas impacted by either one of the events. Compound refers to areas impacted by both drought and hurricane in the same year

$^{*}p < 0.05$; $^{**}p < 0.01$; $^{***}p < 0.001$

**Supplementary Table 2** | Processing of auxiliary datasets.

| Variable | Mechanism | Source resolution | Processing | Data source |
|---|---|---|---|---|
| Elevation | Low-lying, gently-sloped terrain with topographic depressions can increase the risk of flooding and saltwater intrusion[1–5]. | 10 m | Reprojected to 100m with bilinear resampling. | 3DEP DEM[6] |
| Height above nearest drainage (HAND) | | 30 m | Reprojected to 100m with bilinear resampling. | Copernicus HAND[7] |
| Slope | | 100 m | Computed from 100 m elevation. | 3DEP DEM[6] |
| Topographic position index (TPI) | | 100 m | Computed from 100 m elevation. | 3DEP DEM[6] |
| Topographic wetness index (TWI) | | 100 m | Computed from 100 m elevation. | 3DEP DEM[6] |
| Soil texture | Soil characteristics impact hydraulic conductivity, water retention, and legacy impacts from saltwater intrusion[1,2,8,9]. | 800 m | Reprojected to 100m with nearest neighbor resampling. | gridded national soil survey data (SSURGO and STATSGO)[10,11] |
| Soil cation exchange capacity (CEC) | | 800 m | Reprojected to 100m with nearest neighbor resampling. | gridded national soil survey data (SSURGO and STATSGO)[10,11] |
| Hydric soil fraction (Soil hydric) | | 30 m | Reprojected to 100m with bilinear resampling. | SSURGO[10] |
| Distance to flowline | Drainage channels may enhance saltwater loading, but also freshwater dilution[12,13]. | 100 m | Euclidean distance from flowline (natural or artificial). | USGS national hydrography dataset[14] |
| Density of flowline | | 100 m | Density of flowline (natural or artificial) within 1 km. | USGS national hydrography dataset[14] |
| Distance to coast | Factors directly related to saltwater and flooding exposure[5,15]. Trees located close to open water with high | 100 m | Euclidean distance from the coastline. | NOAA medium shoreline dataset[16] |
| Salinity of the closest open | | 0.125° | Satellite derived-sea surface salinity product. Averaged | Copernicus multi-observation |

| | | | | |
|---|---|---|---|---|
| water body ($Salinity_{nearest}$) | salinity may be more likely to transition to experience hypoxia, salt toxicity, and eventually hydraulic failure[1,2,5]. | | (2012-2023) and bilinear extrapolated to fill gaps near land. Reprojected to 100m with bilinear resampling. | global sea surface salinity product[17] |
| Relative sea- and lake-levels | Rising water levels can lead to increased hypoxia, and in the case of sea-level, increased salinization and amplified storm surge effects[5,18]. | Vector | Tidal gauges with at least 8 years of record were retained, each with > 6 months of observations during 2012–2023. We then calculated mean annual water level using seasonally decomposed monthly water level to remove seasonality and sudden anomalies. | NOAA tidal gauges[19] |
| Cumulative windspeed ($Storm_{cumulative}$) | Storm causes windthrow[20], extreme precipitation, and storm surges can push saltwater inland[15,21]. We assumed the saltwater intrusion is proportional to storm intensity and frequency[20]. | 5 km | Windspeed of each hurricane during 2012 – 2023, reconstructed with NOAA storm track HURDAT2[22] using HURRECON[23] at 5 km resolution (parameter: rmw=47[24] and s_par=1.35[23]). Resampled to 100m with bilinear resampling. For random forest driver analysis, we calculated the cumulative sum of windspeed over the years. | NOAA HURDAT2[22] |
| Cumulative severity of | Drought reduces freshwater input into coastal estuary, | 4 km | Aggregated to monthly SPEI. Then, we | GridMET SPEI-1 year[27,28] |

| | | | | |
|---|---|---|---|---|
| drought (Drought$_{cumulative}$) | reduces plant available water and concentrates salt, raising salinity[1,21,25,26]. | | calculated the cumulative SPEI annually of all droughts (i.e. pixels with monthly SPEI < -1). Resampled to 100m with bilinear resampling. For random forest driver analysis, we calculated the cumulative sum of SPEI value over the years. | |
| SPEI anomaly ($\Delta$ SPEI) | | 4 km | Aggregated to monthly SPEI. Then, we calculated mean SPEI (2012-2023) relative to baseline mean (1980-2010). Resampled to 100m with bilinear resampling. | GridMET SPEI-1 year [27,28] |
| Forest type | Flood or salt resistance depends on tree's physiology and forest type[13,21,29]. | 30 m | Reprojected to 100m with nearest neighbor resampling. We also tested other gridded forest type products (e.g. USFS tree species parameter), but the results were similar. | USFS TreeMap 2016[30] |
| Tree cover | Dense canopies compete strongly for essential resources (e.g. freshwater, nutrient)[31]. Higher tree cover may also increase the number of dead trees detected. | 30 m | Resampled to 100m with bilinear resampling. | Hansen tree cover in 2010[32] |

| | | | | |
|---|---|---|---|---|
| Burned area | Fire cause widespread forest loss | 30 m | Resampled to 100m with nearest neighbor. | LANDFIRE[33] and USFS IDS[34] |
| Defoliation | Temporary defoliation caused by insect or pathogens. | Vector | Mapping done by trained experts during annual aerial surveys. Defoliation points and polygons associated with disease or insect. Rasterized to 100m | USFS IDS[34] |
| Infestation | Insect or pathogens tend to target stressed-trees for attack, increasing mortality risk[35]. | Vector | Mapping done by trained experts during annual aerial surveys Mortality points and polygons associated with disease or insect. Rasterized to 100m | USFS IDS[34] |
| Mean annual precipitation (MAP) | Vegetation is sensitive to annual and daily rainfall patterns[36], potentially increase hypoxia under extremely wet conditions[1].9/21/2026 12:36:00 PM | 4 km | Mean total annual precipitation (2012-2023). Resampled to 100m | GridMET[27] |
| Precipitation frequency | | 4 km | Mean precipitation frequency (days with precipitation ≥ 1mm) annually during 2012-2023. Resampled to 100m with bilinear resampling. | GridMET[27] |
| Precipitation frequency anomaly (Δ Precip. frequency) | | 4 km | Mean precipitation frequency (days with precipitation ≥ 1mm) annually during 2012-2023 relative to baseline mean (1980-2010), and divided by baseline standard deviation. sampled to 100m with | GridMET[27] |

| | | | | |
|---|---|---|---|---|
| | | | bilinear resampling. | |
| Precipitation intensity anomaly (Δ Precip. intensity) | | 4 km | Mean precipitation intensity (Mean precipitation on days with precipitation ≥ 1mm) annually during 2012-2023 relative to baseline mean (1980-2010), and divided by baseline standard deviation. Resampled to 100m with bilinear resampling. | GridMET[27] |
| Maximum temperature anomaly (Δ $T_{max}$) | Tree mortality is exacerbated by drought and extreme temperature[35,37]. | 4 km | Mean $T_{max}$ annually during 2012-2023 relative to baseline mean (1980-2010), and divided by baseline standard deviation. Resampled to 100m with bilinear resampling. | GridMET[27] |
| Frost days anomaly (Δ Frost days) | Extreme cold days can induce freezing damage, reduced growth, and frost embolism that can enhance mortality risks[38,39] | 4 km | Mean number of frost days annually during 2012-2023 relative to baseline mean (1980-2010), and divided by baseline standard deviation. Resampled to 100m with bilinear resampling. | GridMET[27] |

**Supplementary Table 3** | Environmental variables in random forest models for explaining mortality hotspots in different flood-prone forest systems.

| Category | Low-lying saltwater | Low-lying freshwater | Inland wetland |
|---|---|---|---|
| Geophysical | Elevation | Elevation | Elevation |
| | Slope | Slope | Slope |
| | TPI | TPI | TPI |
| | TWI | TWI | TWI |
| | HAND | HAND | HAND |
| | Distance to coast | Distance to coast | Distance to coast |
| | Distance to flowline | Distance to flowline | Distance to flowline |
| | Density of flowline | Density of flowline | Density of flowline |
| | Soil texture | Soil texture | Soil texture |
| | Soil hydric | Soil hydric | Soil hydric |
| | Soil CEC | | |
| | Salinity$_{nearest}$ | | |
| Climatic | | MAP | MAP[1] |
| | Precip. frequency | Precip. frequency | Precip. frequency |
| | Δ Precip. frequency | Δ Precip. frequency | Δ Precip. frequency |
| | Δ Precip. intensity | Δ Precip. intensity | Δ Precip. intensity |
| | Δ $T_{max}$ | Δ $T_{max}$ | Δ $T_{max}$ |
| | Δ Frost days | Δ Frost days | Δ Frost days |
| | Δ SPEI | | |
| | Drought$_{cumulative}$ | Drought$_{cumulative}$ | Drought$_{cumulative}$ |
| | Storm$_{cumulative}$ | | |
| Biotic | Tree cover | Tree cover | Tree cover |
| | Forest type | Forest type | Forest type |
| | Infestation frequency | Infestation frequency | Infestation frequency |

Models selected slightly different variables since we excluded highly correlated pairs ($R > 0.7$) and retained only variables that influence tree mortality risk in each specific system. All variables have variance inflation factors (VIF) less than 5.

[1]MAP and Storm$_{cumulative}$ had $R = 0.7$ in inland wetland datasets. We therefore ran separate models containing either one of the two variables (MAP-model and Storm-model) while keeping all other variables unchanged. The result shows that the relative importance of MAP (13%) is much higher than Storm$_{cumulative}$ (4%). The final reported model for inland wetlands thus represents the MAP-model.
